# Integrating *in silico* predictions with an engineered tissue assay identifies Perlecan as an age-perturbed re-quiescence cue for muscle stem cells

**DOI:** 10.1101/2024.07.22.604619

**Authors:** Erik Jacques, Pauline Garcia, Orane Mercier, Yechen Hu, Cyril Degletagne, Jade Ravent, Sidy Fall, Maira P. Almeida, Aaron R. Wheeler, Stephane Angers, Penney M. Gilbert, Fabien Le Grand

## Abstract

Skeletal muscle regeneration is mediated by resident muscle stem cells that produce progeny to repair or recreate muscle. Critical to this function is the ability to transition between states of proliferation and quiescence. This balancing act is disrupted with age, leading to eroded regenerative capacity. Notwistanding, mechanisms by which the regenerating niche directs MuSCs return to the dormant state are largely unknown. Since single-cell RNAseq methods exclude the analysis of multinucleated cells, we generated single-nuclei RNAseq datasets of regenerating muscle to capture the full breadth of myogenic progression. With this, we uncovered new transition states between differentiating myocytes and syncytial multinucleated cells. Using cell communication inference tools, we highlighted receptor-ligand interactions between MuSCs and fusing nuclei. We leveraged a bespoke biomimetic 3D niche that induces MuSC quiescence, to filter the predicted interactions using a Cas9-based functional genomics approach. We found the proteoglycan Perlecan (Hspg2) promotes MuSC re-quiescence. *Hspg2* silencing *in vivo*, during muscle regeneration, induced an aging-like phenotype and perturbed MuSC re-quiescence. Notably, the temporal profile and overall levels of Perlecan were altered with age. Exogenous supplementation with Endorepellin (truncated Perlecan) rescued MuSC decline following aged muscle regeneration, offering a new therapeutic target. Thus, coupling *in silico* predictions with a 3D *in vitro* assay followed by *in vivo* investigations revealed a previously inaccessible window of biology; that spatiotemporal coordination of MuSC re-quiescence is directed by fusing myonuclei.

## Introduction

In healthy, unscathed skeletal muscle, muscle stem cells (MuSCs) are found around the terminally differentiated myofibers in a reversible non-proliferative state termed quiescence (one of three G_0_ states). Quiescence prevents active cycling and exhaustion, and is inextricably linked to potency (i.e., stemness).^1^ For tissue regeneration to proceed, quiescence must be sequentially broken to allow MuSC activation, production of transit amplifying cells, differentiation and self-renewal, and then must be reinstated to maintain tissue healing potency. During regeneration, MuSC quiescence and downstream fate choices are orchestrated by dynamic cell-cell interactions and the ensuing signaling pathways. How quiescence re-entry (re-quiescence) takes place during late-stage regeneration remains unclear. This knowledge gap is essential to fill since disruption of quiescence in aged MuSCs correlates with a decline in both MuSC numbers and regenerative function.^2^

Efforts to study stem cell quiescence and the associated niche interactions have proven challenging in many tissues. This is due to the isolation-induced activation phenomenon and difficulties associated with pinpointing mechanisms *in vivo*.^3–5^ In spite of this, many niche cues have been identified as being essential for maintaining quiescence in uninjured muscle (reviewed by Hung et al. (2023)^6^), particularly from the apical myofiber e.g., Dll4, WNT4, OSM and others.^7–9^ Mind-bomb 1 expression by myofibers, as triggered by the hormonal transition during puberty, promotes quiescence in myogenic populations during postnatal development.^10^ Niche cues driving MuSC re-quiescence at the conclusion of adult myogenesis have proven elusive.

Previous studies (reviewed by Sousa-Victor P, García-Prat & Muñoz-Cánoves (2021)^2^) have indicated that re-quiescence occurs between 5 to 10 days post-injury (DPI) in mice.^11^ During this time-frame, cell signaling is compartmentalized within the skeletal muscle. Pervasive ligands and dynamic receptor expression produce different outcomes according to the sender/receiver cell type (e.g, VEGF) and to the layers of extra-cellular matrix (ECM) surrounding myofibers.^12^ To this latter point, quiescent MuSCs are physically ensheathed between their host myofibers and the surrounding basal lamina. The basal component of the immediate niche is a medley of binding proteins and growth factors that change dynamically throughout regeneration, effectively serving as both a barrier and a medium for cell communication.^13^ Basal lamina is composed of laminins (in particular the 2-1-1 isoform), type IV Collagen, Entactin (nidogen-1) and proteoglycans.^14–17^ This thin and flexible non-cellular sheet is built by connective tissue cells and myogenic cells during myofiber formation (even *in vitro*), but exact contributions to this build-up is still unclear *in vivo*. Moreover, the roles of many basal lamina constituents remain unknown in the context of tissue regeneration.^17–19^

While it was previously considered that MuSCs leave their niche following myofiber injury, intravital imaging has eloquently shown that the basement membrane remains intact even as myofibers undergo necrosis and are cleared during regeneration.^20^ Recently, tissue clearing studies have further demonstrated that MuSCs and their progeny are restrained to their respective silos throughout repair.^21^ Thus, MuSC re-quiescence coincides with nascent myofiber formation and basal lamina remodeling *in vivo* (5-7 DPI)^21^. It is most ostensible that the immediate regenerating niche is involved in directing MuSC return to the dormant state. However, it remains unknown which ligands are provided by fusing nuclei and nascent syncytia during adult myogenesis.

Recent work used single cell RNA-seq to resolve mononucleated cell types and their potential interactions during muscle regeneration.^22,23^ However, these datasets exclude all multinucleated cells, which is crucial missing information as nuclei inside syncytia account for roughly two thirds of muscle tissue nuclear content. As such, single cell profiling cannot infer communications from the regenerating apical niche toward MuSCs. To fill this gap, recent studies delineated the transcriptome of single nuclei in syncytial muscle cells.^24–28^ This work mainly focused on uninjured homeostatic myofibers, necessitating the generation of new datasets to interrogate MuSC/myofiber interactions during adult muscle regeneration.

Herein, we report an approach to identify responsible niche ligands driving MuSC re-quiescence. First, we employed single-nuclei RNAseq (snRNAseq) to infer cell-cell communication *in vivo* between MuSCs and all other myogenic populations, including fusing and nascent myonuclei. We then filtered these interactions through a series of biology-informed gates. Candidate ligands were tested via a functional genomics inspired approach using the mini-IDLE model, a biomimetic skeletal muscle niche that induces quiescence unto activated MuSCs.^29^ *In vivo* validation demonstrated that expression of the basal niche component Perlecan by fusing nuclei is necessary for adequate MuSC dynamics during regeneration. Importantly, we found Perlecan expression to be diminished with age. Resetting Perlecan signaling in old regenerating muscle restored a youthful level of MuSC re-quiescence thereby replenishing the old MuSC niche.

## Results

### The myotube template provides a pro-quiescent niche by non-diffused means

To investigate extracellular cues regulating the critical decision point of re-quiescence in MuSCs, our group leveraged the mini-IDLE model to shed light on possible mechanisms of action. We previously demonstrated that the presence of immature cultured myofibers (myotubes) was an essential component enabling activated MuSCs to re-acquire a quiescent-like phenotype.^29^ Thus, we first sought to determine whether myotube template-derived pro-quiescence cues are predominantly secreted as diffusible factors into the milieu or presented locally. To do this, MuSCs were introduced into a tissue made exclusively of the 3D cellulose-reinforced ECM hydrogel (3D ECM/cellulose), which was co-cultured with a myotube template tissue. This prevented physical interactions between the two, but allowed for communication through the medium (**Figure S1A**). MuSCs added to a myotube template (mini-IDLE) led to a stable Pax7+ population and minimal transient cell cycling at all time-points tested (**Figure S1B-C**), as we previously reported.^29^ By contrast, a high proportion of MuSCs engrafted into 3D ECM/cellulose-alone were Ki67+ at 1, 3, and 7 days post-engraftment (DPE), and we observed a concurrent decline in the Pax7+ population over time (**Figure S1B-C**). MuSCs engrafted in 3D ECM/cellulose and co-cultured with myotube templates showed an initial expansion of the engrafted Pax7+ pool, followed by a later loss of Pax7 expression by most of the daughter cells. By contrast to the 3D ECM/cellulose-only condition, there remained a small population of Pax7+Ki67-cells at 7 DPE (**Figure S1B-C**). Immunohistochemical analysis shows that the co-cultured MuSCs display an intermediate morphology between the elongated mini-IDLE MuSCs and the smaller spherical counterparts in 3D ECM/cellulose-only (**Figure S1D**). Therefore, diffused cues alone are insufficient to phenocopy the quiescent signatures obtained by MuSCs cultured in the mini-IDLE system, which reside in a myotube niche.

To establish a candidate list of putative pro-quiescence drivers, we profiled the myotube template proteome using mass spectrometry. By comparing the conditioned media and the tissue lysate from day 5 myotube templates (i.e. timepoint of MuSC engraftment) and filtering against 3D ECM/cellulose-alone to omit proteins of non-myotube origin, 388 proteins in the conditioned media and 804 in the tissue lysate were identified (**Figure S1E**). Gene ontology (GO) analysis combining cellular compartment, biological processes, and molecular function terms showed the top twenty terms to be generally involved in metabolism (e.g., GO:BP organonitrogen compound metabolomic process, GO:BP amide metabolomic process) and anabolic processes (e.g., GO:BP cytoplasmic translation, GO:BP peptide biosynthetic process, GO:CC contractile fiber), consistent with a developing myotube (**Figure S1F**). Upon closer inspection, there were also proteins associated with cell signaling mediated through ECM and cell junction localized events (**Figure S1G**). These data further argue for the physical proximity of MuSCs to myotubes as a driver of niche induced phenotypic changes.

### Single nuclei RNAseq profiling of young and aged regenerating skeletal muscle reveals the breadth of myogenic trajectory

The next aim was to take a step back and consider/determine/predict/assess how the myogenic populations, analogous to those in mini-IDLE, may be signaling *in vivo* during the period of muscle regeneration when MuSCs re-enter quiescence. Since myocytes and multinucleated cells such as myofibers are poorly represented in scRNAseq datasets, we instead implemented/pursued a single-nuclei RNAseq (snRNAseq) based approach.^30^ Specifically, we isolated and sequenced nuclei isolated from tibialis anterior (TA) muscles of young mice at time-points characterized by progenitor transit amplification and nascent myofiber formation (4 DPI) and by MuSC re-quiescence (7 DPI), which we combined with a previously published 0 DPI snRNAseq dataset (**Figure 1A**; GSE147127^25^). Single nuclei were also collected from the TA muscles of aged mice, at the identical time-points, for snRNAseq data acquisition. After standardized quality control, data wrangling, clustering, manual cell annotated and integration, snRNAseq analysis captured the full myogenic progression for both young and aged skeletal muscle (**Figure 1B-D**). At 0 DPI, the majority of myonuclei split into those associated with fiber sub-types (IIB, IIX and IIA), in addition to the specialized myonuclei of the myotendinous and neuromuscular junctions (MTJ, NMJ), as previously annotated^25^ (**Figure S2**). In contrast, on 4 DPI when transit amplifying cells are produced and new fibers are formed, MuSCs and their cycling counterparts are present (*Pax7*, *Notch3*, *Mki67*, *Lockd*). In addition, we identified myocytes (*Myog*, *Ncam1*) that were undergoing fusion, surmised based upon co-expression of early myosin isoforms, fusion proteins and other fusion essential genes (*Myh3*, *Myh8*, *Jam3*, *Ryr1*, *Camk2a*), as identified by others (**Figures S2, S3**).^24,31–33^ We also noted a non-negligible population of myogenic cells (*Ncam1*) that were enriched for immune-related genes (*C1qb*, *C1qa*, *Ctss*), previously described as “immunomyoblasts” (**Figures S2, S3**).^23^ There were also newly formed myonuclei with reemerging fiber type specifications (*Ttn*, *Myh4*, *Myh1*, *Myh2*) (**Figures S2, S3**). Included as well were immune cells (Macrophages, T cells, B cells, NK cells), fibro-adipogenic progenitors (FAPs), and endothelial cells (ECs). At 7 DPI, similar populations were found with a skew towards increased myonuclei and decreased cycling MuSCs (**Figures S2, S3**). We also noted the appearance of Rian+ myonuclei, as previously reported by others.^25^ This unique dataset, capturing all cell types (including the entire myogenic lineage) in accurate proportions, provided the framework with which to interrogate skeletal muscle inter-cellular signaling.

**Figure 1.**
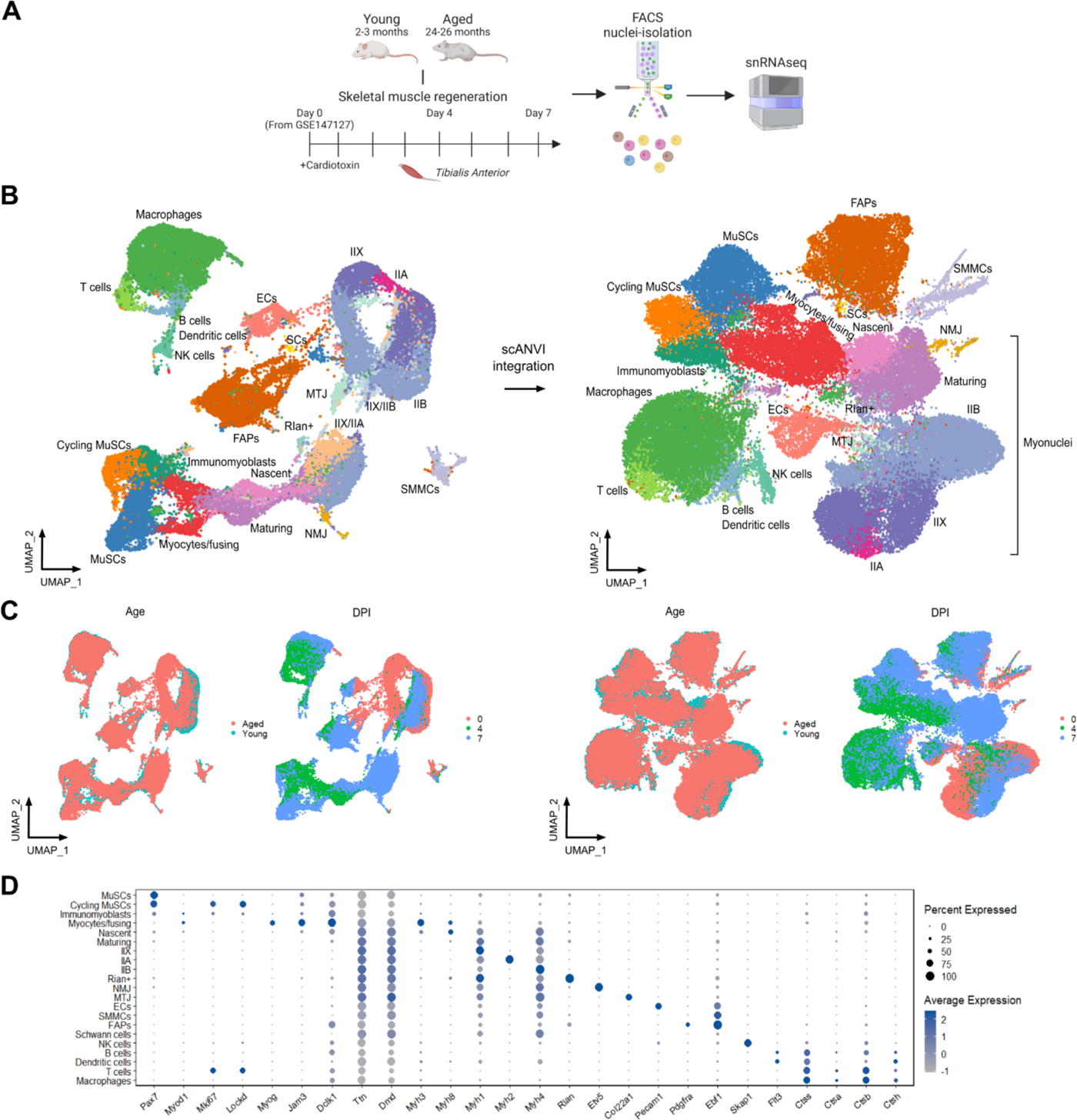
Single-nuclei RNAseq of regenerating muscle. **(A)** Schematic representation of the experimental workflow for snRNAseq of young and aged regenerating skeletal muscle (made with BioRender). N=3 independent biological replicates (different mice) pooled for sequencing. **(B)** Representative UMAP plots of merged 0, 4 and 7 days post injury (DPI) samples from Young and Aged before (left) and after (right) integration. 0 DPI timepoint from GSE147127. **(C)** Representative UMAP plots before (left) and after (right) integration, with color coding according to Age and DPI. **(D)** Dot plot showing selected markers expressed across all identified cell populations. Color and size code for expression level and percent of cells expressed, respectively.

### *Hspg2* and *Lamc1* are candidate pro-quiescence ligands emanating from regenerating myofibers

To predict how MuSCs interact with the other cell populations in their *in vivo* niche, we conducted cell-cell communication inference (CCCI) on the snRNAseq datasets. Using a curated database of receptor-ligand interactions (CellchatDB.mouse)^34^ the transcript counts of one receptor gene in one cell type are combined with the corresponding ligand gene counts in a different cell type to create an interaction strength score. The interactions scores of one or all pathways are then aggregated to look at incoming versus outgoing signal strength between each cell type. We first focused specifically on the snRNAseq dataset produced from the young muscle samples. For this analysis, we implemented the assumption that pro-quiescence cues are not fiber type specific and proceeded to ignore the myonuclear subtypes in favour of a focus on immature as compared to mature myonuclei (based on signature overlap with those from 0 DPI) (**Figure 2A**). Afterwards, the total incoming and outgoing interaction strengths of each cell type were split by timepoint for a broad overview of muscle signaling (**Figure 2B**). This showed that, as expected, at rest (0 DPI) the mature myonuclei (myofibres) are the primary communicators, making up over 90% of the nuclei in the tissue.^35^ At 4 DPI, myocytes are the major players along with MuSCs, in addition to the macrophages and FAPs that are clearing away debris and rebuilding the interstitium, respectively. By 7 DPI, a timepoint highlighted by others as the transition point for MuSCs re-quiescence^11^, MuSCs now become the primary communication hub for both incoming and outgoing interaction strength. Another prominent, yet previously unreported player, are the newly formed, yet immature myonuclei (**Figure 2B**). With a closer look at myogenic cell signaling pathways, we notice that at rest, MuSCs are the recipients of well-known Notch, Laminin, Ncam activity, in addition to Sema6, Hspg, Fgf and Bmp (**Figure 2C**). Myofibers show activated growth factor signaling (Fgf, Bmp, Vegf), ECM signaling (Laminin, Hspg), and Visfatin signaling, which is involved in calcium homeostasis^36^. At 4 DPI, myocytes emerge prominently, particularly with regards to incoming signal strength, and are involved in a plethora of pathways such as Laminin, Fn1, Bmp, Igf, etc, all originating from FAPs. MuSCs on the other hand, display sustained Notch and Sema6 activity, in addition to Hgf, Egf, PI3K, to name a few (**Figure 2D**). Similar signaling pathways, albeit with increased or decreased activity, preside over 7 DPI, as well as new pathways including Wnt, ncWnt, and Cdh (**Figure 2E**).

**Figure 2.**
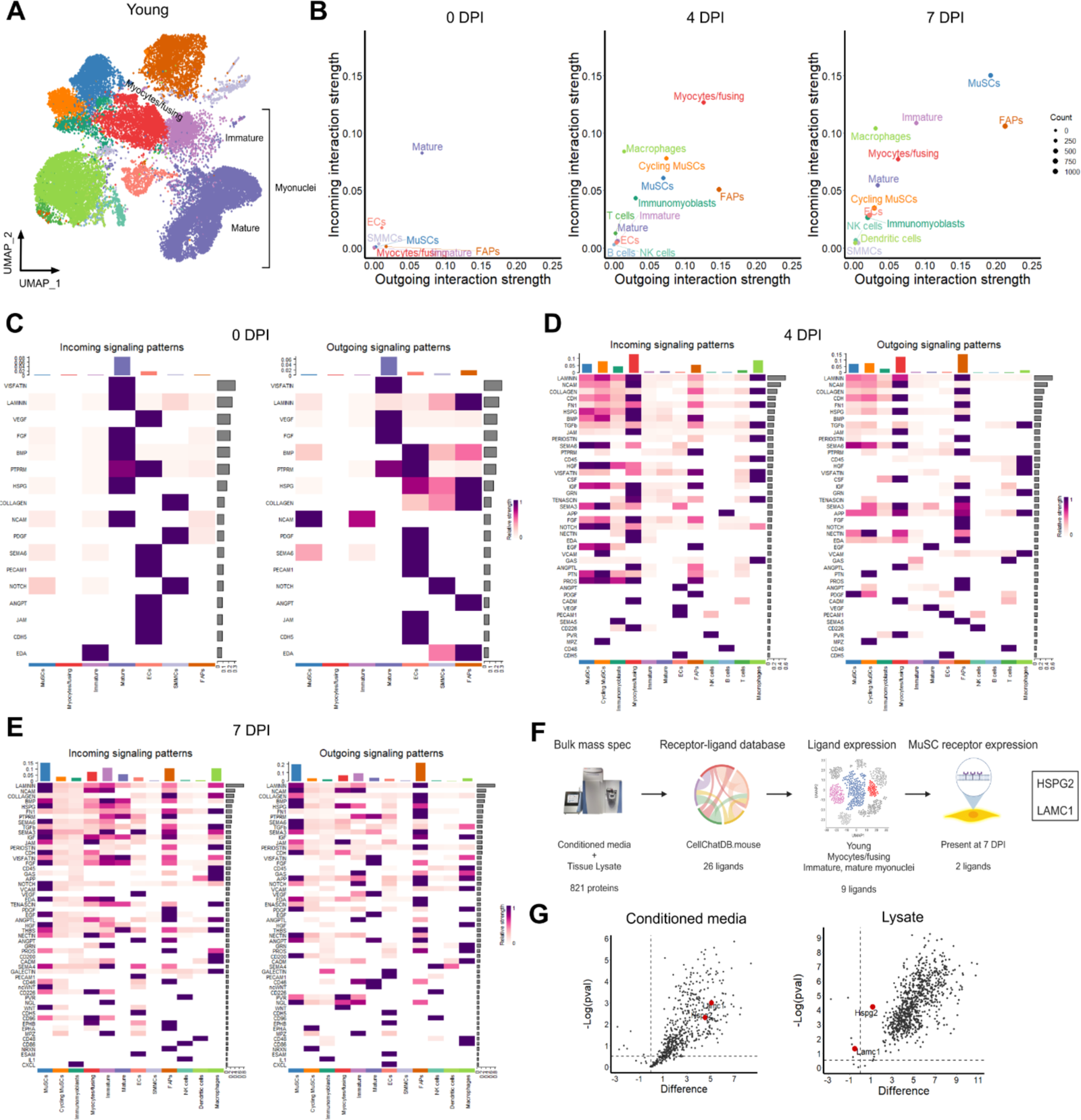
Cell-cell ligand-receptor inference shows MuSC communication and shortlists candidate pro-quiescent cues. **(A)** Representative UMAP plot of the Young data subset with the myonuclei highlighted and split into “Immature” and “Mature”. **(B)** Dot plot showing the overall outgoing and incoming interaction strengths (as calculated by CellChat) for each cell type across each day post injury (DPI) timepoint. Dot size codes for the number of interactions (count) for each cell type at that timepoint. **(C-E)** Heatmaps showing the relative strength of incoming and outgoing signalling pathways per cell type for 0 (C), 4 (D), and 7 DPI (E). Total signal strength for each pathway and cell type are summed on the second x-axis (top, color-coded) and y-axis (right, greyscale) of each heatmap. **(F)** Schematic representation of the filtering steps for candidate ligand selection (made with BioRender). **(G)** Dot plot of the proteins identified by mass spectrometry in the conditioned media (left) and myotube template lysate (right) plotted by the Difference and -Log(pval) on the x and y-axis. Values were calculated by comparing against ECM-only tissue samples as controls. Candidates *Hspg2* and *Lamc1* are highlighted in with red circles. n=3 tissue replicates and results were aggregated together for each experimental condition.

To select candidate cues among a medley of pathways where it is suspected that those originating from myogenic populations mediate MuSC re-quiescence, we employed a series of logic-gates that combined the mini-IDLE and snRNAseq data (**Figure 2F)**. The 821 proteins identified by bulk mass spectrometry were filtered to remove those that were absent from the CellChatDB,mouse curated protein ligand list, which led to 26 total ligands. These were filtered once again by a requirement for expression by any of the following niche cell populations: myocytes/fusing, immature, and mature myonuclei. Finally, with the 9 remaining ligands, we shortlisted those for which the ligands plus the corresponding MuSC receptors were expressed at 7 DPI, with the speculation that signals received here would be involved in inducing quiescence. The ligand genes that met all criteria were *Hspg2* (Heperan sulfate proteoglycan 2) and *Lamc1* (Laminin subunit gamma 1) (**Figure 2F**). The CCCI data shows Laminin and Hspg pathway activity are present in MuSCs at 0 DPI and ramps up in activity from 4 to 7 DPI, to eventually return to pre-injury levels (**Figure 2C-E**). Both HSPG2 and LAMC1 were identified in the mini-IDLE myotube template conditioned media, indicating that myotubes actively produce both proteins (**Figure 2G**). Only HSPG2 was found in the myotube template tissue lysate, suggesting that this ECM protein was being physically deposited into the tissue. Conversely, LAMC1 was found at slightly higher levels in the control samples (ECM-only). This suggests that LAMC1 is either not being deposited into the ECM, or that the levels of this subunit being produced are minimal compared to that contributed by the Geltrex® incorporated into the 3D ECM base (**Figure 2G**).

### Myotube-derived *Hspg2* is essential for MuSC quiescence in mini-IDLE

Inspired by functional genomic screens, we next used a CRISPR/Cas9 approach to knock-down the candidate ligand genes in the myotube template and then assessed samples to determine whether perturbations disrupted MuSC re-quiescence in mini-IDLE (using multi-variate image-based readouts for assessing MuSC phenotype). A Cas9 myoblast line was created by harvesting hindlimb muscle tissue from the Gt(ROSA)26Sortm1.1(CAG-cas9*,-EGFP)Fezh/J mouse, which in turn was used to create Cas9 expressing myotube templates. In parallel, two single-guide RNAs (sgRNAs) were designed per gene for perturbation studies. Sequences were ligated to the pLCKO plasmid, cloned, and packaged for lentiviral delivery. sgRNAs were also designed for a set of positive controls; genes for which disorder would knowingly alter the MuSC phenotype. Targeting *Psmb2*, an essential gene and member of the proteosome^37^, would lead to cell death, mimicking muscle tissue injury. *Osm* (Oncostatin-M) and *Cdh15* (M-cadherin), are each produced by the myotube template (**Figure S4A**) and are described for their roles in maintaining quiescence *in vivo*.^38,39^ Packaged sgRNAs were tested in a 2D setting on both myoblasts and myotubes formed using the Cas9 line, which revealed an average editing efficiency of ≈80% across all experimental conditions (**Figure S4B-D**). We then established the viral load by testing varying amounts of the packaged sgRNAs and moved forward with the viral load in which *Psmb2* sgRNAs, but not pLCKO sgRNAs, altered myotube viability (**Figure S4E,F**) and MuSC phenotype (**Figure S4G,H**).

We evaluated two lentiviral delivery regimes were in the course of screening the candidate ligands in mini-IDLE: (a) Day -1 (24 hours post-seeding), to perturb proteins that are suspected of being deposited into the ECM over time, with the caveat of potentially altering myotube formation; (b) and Day 4, such that myotubes are formed before editing, with the caveat that editing may be less efficient in multinucleated cells (**Figure 3A**). Myotube template total knockdowns (KDs) were generated, YFP+ MuSCs were engrafted and then analysed at the 7 DPE endpoint across the following previously established metrics^29^: Pax7+ donor cell enumeration; percentage MyoD+; Mean donor cell elongation and nuclear eccentricity; and total YFP+ tissue coverage (**Figure 3A**). Distilling the results into principal components revealed a meta-cluster composed of control, pLCKO, and *Lamc1*; with separate clusters for *Osm*, *Cdh15*, *Psmb2* and *Hspg2* (Day -1) (**Figure 3B**). Histologically, *Psmb2* sgRNAs caused reduced and poor quality myotubes (**Figure S5**), which translated to activation (MyoD+) and differentiation (increased YFP+ tissue coverage) of engrafted MuSCs (**Figure S6**). Proxy measures of effect size via the mahalanobis distance and Hotelling’s T-square test showed a significant overall perturbation of the MuSC phenotype whether editing at the myoblast or myotube stage (**Figure 3C**). *Osm* sgRNAs, targeting production of the secreted protein Oncostatin-M, showed increased variability along principal component 2 (**Figure 3B**), but similar overall effect sizes (**Figure 3C**), which was explained by alterations in MuSC activation state and morphology (**Figure S6**). *Cdh15* gRNAs also showed enhanced effect sizes manifesting as elevated MyoD activation and alterations in nuclear morphology (**Figures 3C,S6**). Impacts were more prominent when myoblasts were edited on Day -1 which may be due to low protein turnover or a potential confounding variable since myoblast fusion and myotube maturity were each found to be impaired in this group (**Figure S5**). *Lamc1* gRNAs showed no significant alterations (**Figures 3B-C, S5**). Notably, *Hspg2* gRNAs introduced on Day -1 induced significant perturbations to the MuSC phenotype, across nearly all metrics tested, indicating a poor ability to quiescence in min-IDLE tissues when this candidate hit is disrupted (**Figures 3C, S6**).

**Figure 3.**
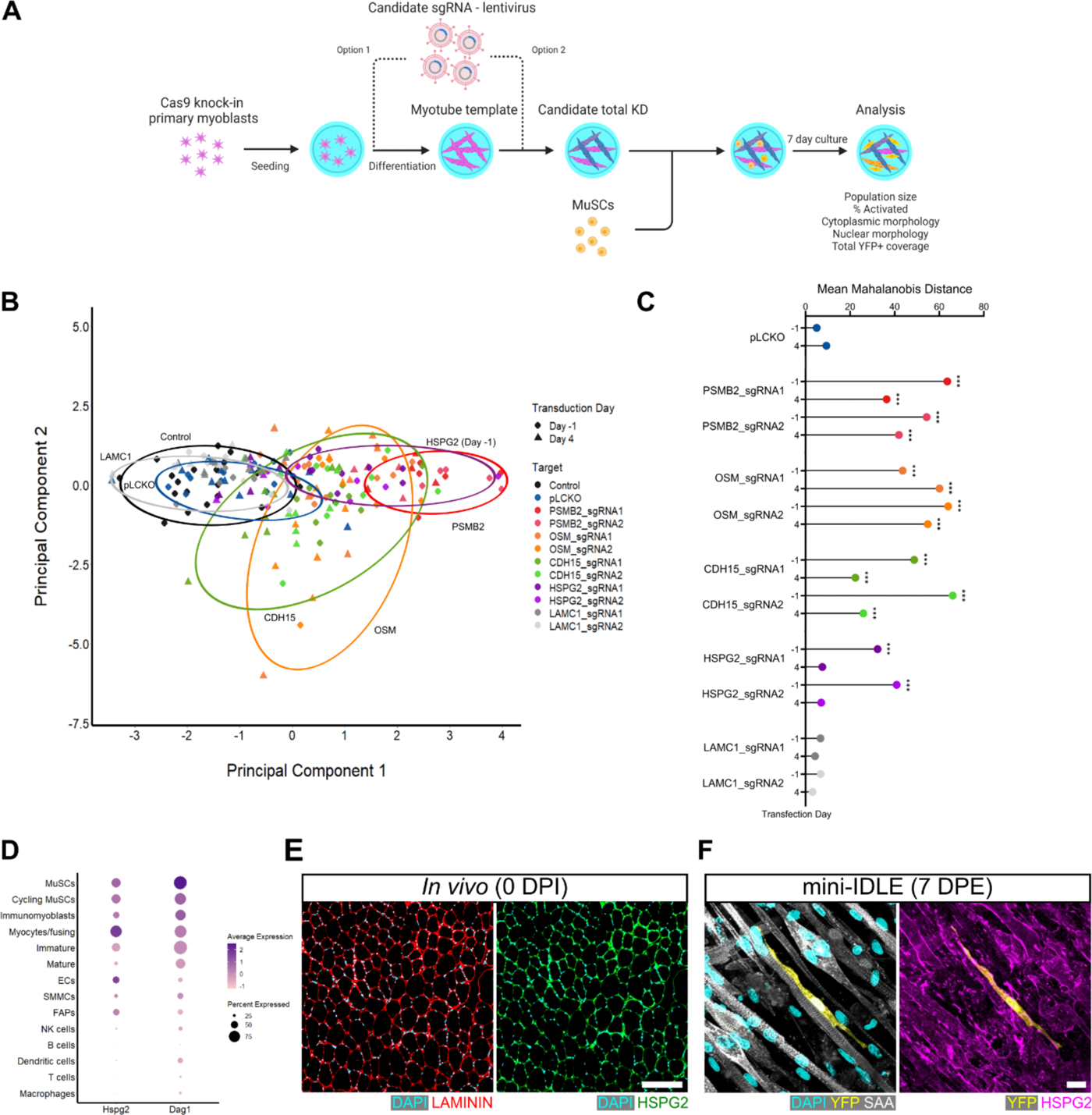
Cas9-mediated screen reveals *Hspg2* is essential for re-quiescence in mini-IDLE. **(A)** Schematic of the experimental setup for Cas9 editing of selected genes in myotube templates and subsequent analysis of engrafted MuSCs (made with BioRender). **(B)** Principal component analysis plot showing the principal component 1 and 2 values for the average MuSC population features of every tissue analysed. Color and shape code for the target site/sgRNA and day of transduction, along with lassos for groups that cluster with, or separate from controls (which center around their respectively centroids). N=4-20 across N=2-6 independent biological replicates. **(C)** Lollipop plot showing the mean Mahalanobis distance for each experimental group compared to the control centroid. N=6-20 across N=3 independent biological replicates; p-values were calculated for Hotelling’s T-squared statistic, ***p˂0.001. **(D)** Dot plot showing gene expression of *Hspg2* and the receptor *Dag1* across all identified cell populations in the Young snRNAseq subset. Color and size code for average expression level and percent of cells expressed, respectively. (**E)** Representative confocal image of perlecan protein (green) *in vivo* in uninjured muscle and also labelled for DAPI (cyan) and laminin (red). Scale bar, 75µm. **(F)** Representative confocal image of perlecan protein (magenta) in a myotube template at 7 DPE engrafted with adult MuSCs (YFP) and also labelled for DAPI (cyan) and sarcomeric α-actinin (SAA, white). Scale bar, 50µm.

Heparan sulfate proteoglycan 2 (HSPG2), also known as Perlecan, is a massive ECM protein modifiable with multiple chains of glycosaminoglycans. The snRNAseq data indicates that Perlecan expression is ramped up during myogenesis and peaks during myocyte fusion/myonuclei formation (**Figure 3D**). The corresponding receptor Dag1 (dystroglycan-1 or α-dystroglycan, a member of the dystroglycan glycoprotein complex) is expressed by MuSCs (**Figure 3D**). Confocal imaging *in vivo* shows Perlecan deposited in the basal lamina (**Figure 3E**). In mini-IDLE, Perlecan is deposited into the ECM and coats the outside of the myotubes, and by 7 DPE, myotube-associated MuSCs are also in close proximity to the protein (**Figure 3F**). All in all, genetic perturbation of select ligand genes within the myotube template showed Perlecan is an essential component to the MuSC phenotype in mini-IDLE.

### *Hspg2* knockdown disrupts myogenesis and impairs MuSC re-quiescence *in vivo*

We next tested the role of Perlecan in muscle regeneration *in vivo* using a targeted but transient siRNA-based knockdown approach, given its temporally restricted production. After a cardiotoxin induced injury to the TA muscle of young mice, control (siCtl) or *Hspg2* (si*Hspg2*) targeting siRNA was injected at 2 and 3 DPI for a 40-48hr downregulation during myogenic differentiation and myofiber formation (**Figure 4A**). To validate the siRNA knock-down strategy *in vivo*, TA muscles harvested at 4 days post-injury (DPI). Cryosections were used as material for qPCR analysis and were also stained to visualize Perlecan, which was quantified as Mean Fluorescence Intensity (MFI). Decreased HSPG2 protein and transcript expression was observed upon analysis of the si*Hspg2* samples when compared to the siCtl (**Figure S7A-D**). TA muscles harvested at 7 DPI exhibited defects in the muscle regeneration process. Small, disorganized myofibers were found at the injury site, with numerous interstitial nuclei observed upon Hematoxylin and Eosin staining (**Figure 4B**). Cross-sectional area quantification confirmed a smaller fiber diameter in the si*Hspg2* condition compared to siCtl (**Figure 4C,D**). There was also extensive fibrosis observed in si*Hspg2* cryosections (**Figure 4H** and **Figure S7E**). Interestingly, at this analysis timepoint there was an increase in the number of nuclei/fiber (**Figure S7E,F**), and many instances of multiple fibers forming within a single larger basal lamina circumference i.e., fibers within fibers (**Figure S7E,G**). To our knowledge this is a unique phenotype never previously reported. No significant differences were observed in embryonic myosin heavy chain staining coupled with adult fast myosin heavy chains (MY-32) staining across the conditions, indicating that defects in myofiber size are not likely caused by developmental stalling (**Figure S7E,H**). Furthermore, there were no overt differences in macrophage content to suggest alterations in debris clearance or inflammation (**Figure S7E,I**). There are thus more myofibers being formed at 7 DPI following a transient *Hspg2* knockdown on DPI 2 and 3.

**Figure 4.**
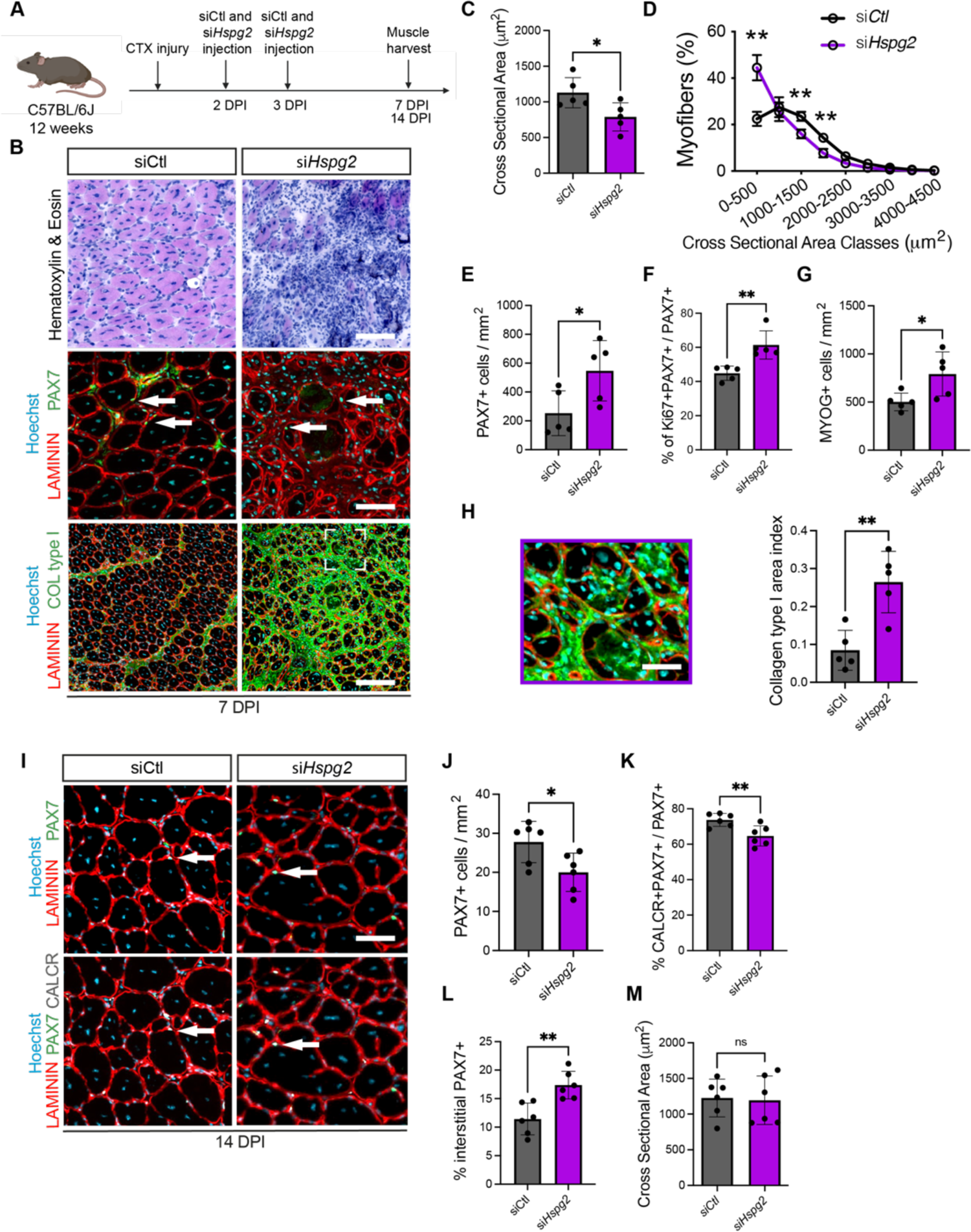
Hspg2 contributes to MuSC re-quiescence during muscle regeneration. **(A)** Experimental scheme of timeline for regeneration studies wherein C57BL6/J mice were injected with cardiotoxin in the Tibialis Anterior (TA) muscle and subsequently injected with siRNA Control (siCtl) or Hspg2 (si*Hspg2*). TA muscles were harvested at 7 and 14 days post-injury (DPI). **(B)** (top) Hematoxylin & Eosin staining, (middle) Pax7 (green) and laminin (red) and (bottom) collagen type I (green) and laminin (red) staining with Hoechst (blue) counterstain performed on 7 DPI TA muscles treated with siCtl (left column) and si*Hspg2* (right column). N=5 mice. Arrows highlight Pax7+ cells. **(C)** Bar graph displaying mean myofiber cross sectional area for CTX-injured TA muscles of siCtl and si*Hspg2* treatment conditions at 7 DPI. N=5 mice. **(D)** Distribution of myofiber cross-sectional area for samples described in C. N=5 mice. **(E)** Quantification of the number of Pax7+ cells per mm^2^ on cryosections at 7 DPI. N=5 mice. **(F)** Quantification of the number Pax7+Ki67+/Pax7+ in sections from siCtl and si*Hspg2* treated muscles at 7 DPI. N=5 mice. **(G)** Quantification of Myog+ cells in sections from siCtl and si*Hspg2* treated muscles at 7 DPI. N=5 mice. **(H)** Representative image (left) and quantification (right) of Collagen type I area in sections from siCtl and si*Hspg2* treated muscles at 7 DPI. N=5 mice. **(I)** Representative images of calcitonin receptor (CalcR, gray), Pax7 (green) and Laminin (red) immunostaining and Hoechst counterstain (blue) in sections from siCtl and siHspg2 treatment group muscles after 14 DPI. N=6 mice. **(J-L)** Bar graph quantification of the number of **(J)** PAX7+ cells, **(K)** CalcR+Pax7+ cells, and **(L)** Pax7+ cells located in the interstitial space per mm^2^ on cryosections from siCtl and si*Hspg2* treated muscles at 14 DPI. N=6 mice. **(M)** Bar graph displaying mean myofiber cross sectional area for CTX-injured TA muscles of siCtl and si*Hspg2* treatment conditions at 14 DPI. N=6 mice. p-values were calculated using the Student’s t-test, **p˂0.005, *p˂0.05, ns: non significative. Scale bars (B) 50 μm, 30 μm, 200 μm, (H) 25 μm (I) 40 μm.

Looking specifically at the MuSC compartment (**Figure 4B**), we quantified a near three-fold increase in population size at this timepoint (**Figure 4E**). These were not quiescent (CalcR-), but rather an increased proportion in cell cycle (Ki67+) (**Figure 4F** and **Figure S7J,K**). Additionally, we quantified a near two-fold increase in the number of terminally differentiated myocytes (**Figure 4G**). That being so, at 7 DPI there was a surplus of myogenic cells. Looking out to 14 DPI, when the MuSC pool is converting to pre-injury status, there was now a reduced number of MuSCs within the si*Hspg2* condition (**Figure 4I,J**), and a smaller proportion were found to be quiescent (CalcR+) (**Figure 4I,K**), with a concomitant increase in the proportion of MuSCs in the interstitial space i.e., the opposite side of the basal lamina (**Figure 4L**). Meanwhile, we found that myofiber cross-sectional area increased to be comparable again siCtl at this timepoint, though there was a loss of type IIA and a gain of type IIB myofibers (**Figure 4M** and **Figure S7L**). Altogether, *Hspg2* knockdown during early regeneration led to an overproduction of myogenic cells and a reduction in the quiescent MuSC pool after myofiber formation. These results provide evidence of the crucial role of *Hspg2* during muscle regeneration. The significant consequences of *Hspg2* loss on MuSCs corroborate our mini-IDLE results of the former being essential to re-quiescence.

### Exogenous addition of Endorepellin blunts unruly MuSC activation

We next sought to demonstrate that Endorepellin reconstitution in *Hspg2* depleted muscle environments can rescue MuSC re-quiescence. As an alternative to full-length Perlecan, which is not sold recombinantly, we investigated recombinant Endorepellin, a truncated version of the Perlecan domain V that has been shown to act as a bioactive signaling ligand^40^ and includes the Dag1 binding region. First, we treated freshly sorted MuSCs with Endorepellin over the first 72-hours of culture (a period of activation) and showed that Endorepellin treatment delays MuSC activation and proliferation (**Figure S8A-C**). Next, to understand the contribution of Endorepellin to the quiescent MuSC phenotype in mini-IDLE, Endorepellin was added to the assay in the context of *Hspg2* total knock-down (**Figure 5A**). Strikingly, this bioactive ligand was able to push the MuSC population back towards a quiescent-like phenotype more similar to control (**Figure 5B**). Looking in greater detail, trends indicate that a rescue in the population size and MuSC activation state prevented precocious differentiation; however, nuclear but not cytoplasmic morphology was rescued (**Figure S8D-H**). We next sought to rescue the *in vivo Hspg2* siRNA phenotype, and pursued experiments where siCtl or si*Hspg2*, each supplemented with Endorepellin, were conducted in young C57BL/6J mice (**Figure 5C**). A restoration of MuSC number was observed at 7 DPI, and was associated with a decrease in the proportion of proliferating MuSCs in the Endorepellin-supplemented condition (**Figures 5D, E, F**). Additionally, a reduction in differentiated cells was observed (**Figure 5G**). All in all, these studies demonstrate that Endorepellin, a bioactive ligand and proxy for full-length Perlecan, rescues most of the *in vitro* and *in vivo* phenotypes we observed in MuSCs following *Hspg2* disruption in the niche.

**Figure 5.**
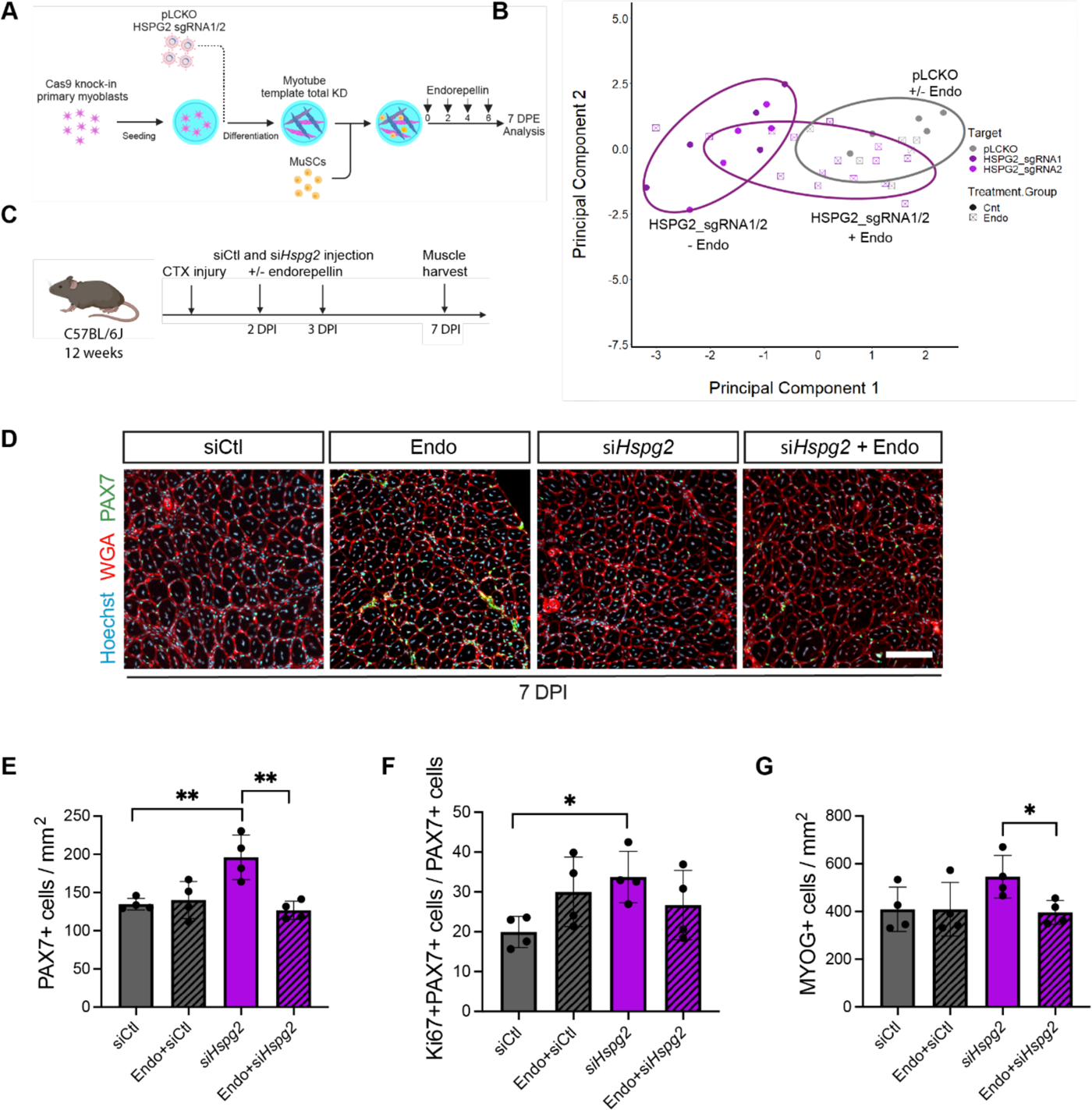
*Hspg2* loss is partially rescued by Endorepellin treatment. **(A)** Schematic of the experimental setup for the rescue of *Hspg2* knockdown in mini-IDLE with Endorepellin (made with BioRender). **(B)** Principal component analysis plot showing the principal component 1 and 2 values for the average MuSC population features of every tissue analyzed. Color and shape code for the target site/sgRNA and treatment group (control versus Endorepellin), along with lassos which center around their respective centroids. n=5-8 tissues across N=2-3 independent biological replicates. **(C)** Experimental scheme for the in vivo rescue of *Hspg2* knockdown with Endorepellin (made with BioRender). **(D)** Representative Pax7 immunostaining in siControl (siCtl), siCtl+Endorepellin, si*Hspg2* and si*Hspg2*+Endorepellin muscles at 7 days post-injury (DPI). N=4 mice. **(E-G)** Bar graphs displaying quantification of the number of **(E)** Pax7+ cells, **(F)** Pax7+Ki67+ cells, **(G)** Myog+ cells per mm^2^ in siCtl, si*Hspg2*, siCtl+Endorepellin and si*Hspg2*+Endorepellin treatment conditions at 7 DPI. N=4 mice. Plots display mean ± s.d. with individual biological replicates (n=4 mice); one-way ANOVA with Tukey’s post-test. **p˂0.005, *p˂0.05. Scale bars (D) 100 μm.

### Reduced Perlecan signaling is causal to age-induced erosion of MuSC re-quiescence

Having identified an essential cue in the regenerating niche for MuSC re-quiescence, we next interrogated whether it is disrupted with age, where MuSCs are reduced in number and in a ‘shallower’ state of quiescence according to -omic signatures. CCCI comparisons between the young and aged snRNAseq datasets show that there is an over two-fold reduction in communication probability in the aged, from both the myocytes/fusing and immature myonuclei (**Figure 6A**). Next asking whether the ligand, receptor, or both are dysregulated; differential expression revealed that *Hspg2* is downregulated in the myocytes, and even more so in the myonuclei, suggesting that Perlecan production is reduced and shorter-lived in the myogenic program (**Figure 6B**). Moreover, *Dag1* receptor expression is also downregulated in aged MuSCs (**Figure 6B**). To confirm Perlecan dysregulation at the protein level, muscle tissue lysate was collected from young and aged mice at various timepoints after injury. Western blots reveal that Perlecan in young muscle begins at low levels and peaks at nearly eight-fold at 4 DPI (**Figure 6C-D**). From 4-14 DPI, protein levels trend back to baseline over the course of ECM remodeling (**Figure 6C-D**). In the aged muscle, baseline Perlecan levels are over three-fold lower (**Figure 6C,E**). Production is also severely impaired at 4 DPI and the peak is delayed to 7 DPI, though still at lower levels than observed in young muscle (**Figure 6C,E**). We also compared another component of the basal lamina: laminin (α-2 subunit). Laminin also starts at low levels and peaks at over eight-fold by 4 DPI (**Figure S9A-B**). In contrast to Perlecan, it remains relatively stable until 14 DPI and does not yet trend back to baseline (**Figure S9A-B**). In the aged muscle, baseline is two-fold lower, production is nearly nil at 4 DPI, and yet delayed to 7 DPI at levels comparable to young muscle 4 DPI (**Figure S9A,C**).

**Figure 6.**
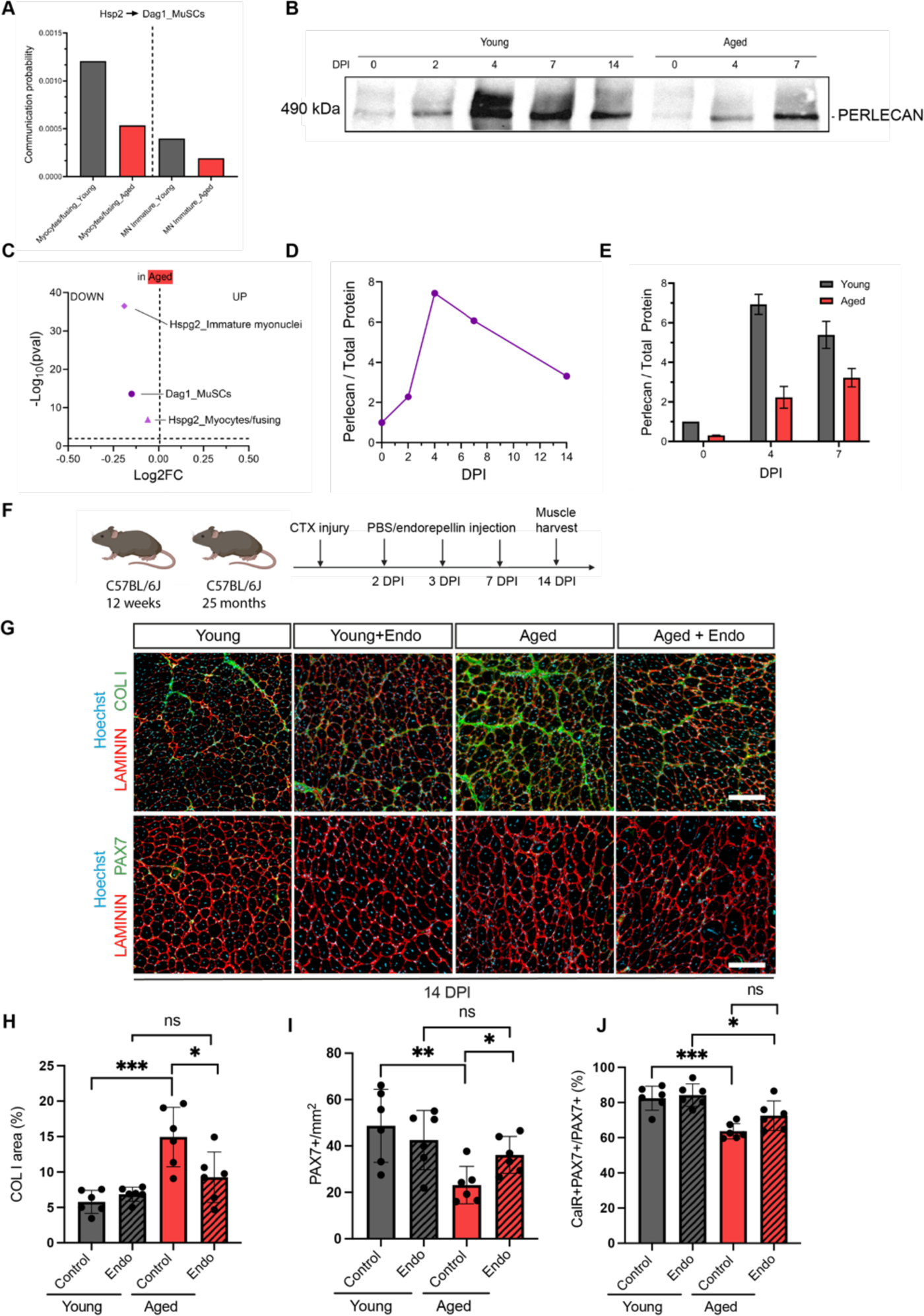
Endorepellin rescues MuSC re-quiescence in aged muscle. **(A)** Bar plot showing the calculated communication probability between *Hspg2* (myocytes or immature myonuclei) and *Dag1* from the MuSCs in young (grey) and aged (red) samples. **(B)** Representative western blot for perlecan in the young vs aged across days post-injury (DPI). **(C)** Volcano plot showing *Hspg2* (in myocytes and immature myonuclei) and *Dag1* (in MuSCs) log_2_ fold-change (FC) in the aged. **(D)** Line graph of perlecan protein quantification in young muscle samples across DPI. **(E)** Bar plot showing mean ± s.e.m. perlecan levels in the young (red) and aged (grey) samples at 0, 4 and 7 DPI. **(F)** Schematic of timeline for in vivo aged muscle Endorepellin rescue experiments. **(G)** Representative images of Collagen I, Pax7 and Laminin immunostaining in muscle sections from young and aged mice supplemented with Endorepellin 14 days after cardiotoxin injury. Scale bar 150 μm **(H-J)** Quantification of **(H)** Collagen I area, **(I)** Pax7+ cells in young, young+Endorepellin, aged, and aged+Endorepellin mice condition (n=6 mice). **(I)** Quantification of Pax7+ cells and **(J)** CalcR+ Pax7+ cells in muscle sections analyzed from young, young+Endorepellin, aged, and aged+Endorepellin treatment conditions. N=6 mice. Plots display mean ± s.d. with individual biological replicates (n=6 mice); one-way ANOVA with Tukey’s post-test. ***p˂0.001, **p˂0.005, *p˂0.05, ns: non-significant.

With a loss in Perlecan signaling potentially explaining the poor return to quiescence that characterizes aged MuSCs after regeneration, we next asked whether Endorepellin could serve as a rescue ligand. Local injections at 2, 3 and 7 DPI were administered to young and aged muscle and then analyzed at 14 DPI where MuSCs are normally found quiescent (**Figure 6F**). Interestingly, quantification of the collagen I area demonstrated that aged mice supplemented with Endorepellin showed a decreased area of fibrosis compared to untreated aged mice (and opposite to si*Hspg2*) (**Figure 6G, 6H**). However, myofiber CSA was not rescued, with increased proportion of smaller myofibers observed in aged muscle compared to young muscle (**Figure S9D**). Even so, Pax7+ numbers, reduced nearly three-fold at this timepoint in aged muscle, were partially rescued two-fold with the Endorepellin treatment (**Figure 6G,I**). Finally, we explored MuSC quiescence using the CalcR marker and observed an increase in the proportion of quiescent MuSCs in aged mice supplemented with Endorepellin (**Figure 6J**). Thus, Endorepellin supplementation in the regenerating niche was able to rescue the quiescence-status of the aged MuSC population.

## Discussion

We report the discovery of Perlecan as an essential protein dictating MuSC re-quiescence in the regenerating muscle niche. We utilized snRNAseq to get one of the first looks at gene expression signatures across all myogenic cells during adult myogenesis since mini-IDLE data indicated the requirement of the myotube component to drive re-quiescence. This snRNAseq dataset, a critical resource for our study and others in the future, captures global cell-cell communication in skeletal muscle. With this, we found that regeneration time-points associated with renewal of the MuSC population are characterized by extensive signaling. Using a bioinformatics-based approach, refined by logic filters and integrated with the phenotypic assessment made possible by the mini- IDLE assay, we reveal a process by which fusing myonuclei contribute to a Perlecan-based remodeling of the basal lamina to drive MuSC re-quiescence. Restoring Perlecan signaling in the context of aged muscle rescues self-renewal to maintain the aged MuSC population and offers a new mechanism driving MuSC exhaustion over the lifespan.

An outstanding question proposed by the field is what instructs myogenic cell differentiation or self-renewal?^41^ Our data indicate that they operate hand-in-hand. Since the advent of omics signatures, it’s been clear that myogenic cells construct their ECM.^42,43^ As these proteins are deposited in the immediate vicinity, studies have contributed to a mechanistic understanding of how MuSCs specifically contribute, and how these components influence cell functions. E.g., under Notch signaling, MuSCs integrate with the basal lamina and contribute proteins such as collagen V to maintain quiescence; upon activation, fibronectin and laminin 1-1-1 promote expansion.^43–46^ The temporally restricted production of Perlecan in fusing myocytes/nascent myonuclei suggests a uniqueness to the ECM produced later into the myogenic program and leaves much to be discovered about basal lamina remodeling. Additionally, our observations indicate that differentiation occurs concurrently with self-renewal, with an unforeseen ‘backwards’ communication from committed to uncommitted cells.

The use of mini-IDLE demonstrates the advantages of *in vitro* engineered tissues applied to the study of MuSC biology. Independent manipulation of the distinct tissue compartments (myotube, ECM, MuSCs, medium) allowed for the luring of niche cues and a new method for bioinformatics derived candidate screening. The determination that physical association between myotubes and MuSCs led to our targeted candidate selection. Moreover, Cas9-mediated perturbation allowed for the assessment of individual gene contributions to the MuSC phenotype. *Psmb2* sgRNAs led to poor myotube formation and viability which presumably served as an injury mimic to MuSCs. We also selected to known cues for quiescence maintenance: *Cdh15* and *Osm*. *Cdh15* (or M-cadherin, which forms adherence junction with other M-cadherins) is expressed in MuSCs and myofibers, along with *Cdh2* (N-cadherin). Intriguingly, *Cdh15* knockout in MuSCs *in vivo* produced a minimal phenotype which was compounded during a *Cdh2* double knockout.^39^ Here perturbation of *Cdh15* in the myotubes led to a similar effect size, affecting MuSC morphology and activation state; which may also be impacted by non-ubiquitous cadherin expression in mini-IDLE MuSCs.^29^ Effects were more pronounced when perturbing on Day -1 as this impaired myotube formation as well.^47,48^ Conversely, *Osm* perturbation did not affect the myotubes, as expected (though elevated level induce atrophy), and produced strong effects on the MuSCs, indicating that a secreted inducer being removed (despite Perlecan being present) can still have effects (discussed below).^38,49^ Then, sgRNAs for our candidates produced various results. *Lamc1* perturbation did not lead to any changes in myotubes or MuSCs despite high editing efficiency. While *Lamc1* whole knockout mice demonstrate embryonic lethality^50^ and laminins are a primary component of the quiescent MuSC ECM, this γ-1 subunit may not specifically mediate quiescence signaling (though it is upregulated with injury ^46^) or impair laminin fibrillogenesis due to compensatory mechanisms. Moreover, it may be due to a technical limitation related to addition of Geltrex©. Our mass spec data indicates that *Lamc1* was not preferentially identified in the lysate compared to control, meaning this protein was likely in the ECM, which could have masked a Cas9 disruption in the myotubes. Here again however, the modularity of such models would allow future studies to circumvent such a limitation by utilizing different hydrogel formulations. Lastly, *Hspg2* perturbation produced a hit exclusively on Day -1 when preventing buildup in the ECM. Notably, the MuSC response was differentiation, similar to 2D cultures that are also absent quiescence cues.

Proteoglycans represent a yet unexplored subcomponent of the basal lamina. Perlecan (HSPG2), named for its pearls-on-a-string appearance under EM imaging, is a large ≈500 kDA protein with glycoaminoglycan chains (GAGs). It is a basement membrane building block containing receptor binding sites (for integrins and dystroglycan), and is also a major binder of growth factors.^16,51–54^ Along with other PGs, it interconnects the laminin and collagen IV networks.^46,55^ First identified in skeletal muscle just before the turn of the century, Perlecan interacts with the myofiber sarcolemma via glycosylphosphatidylinositol anchors.^56–58^ What’s more, GAG chains and their sulfation was demonstrated as important for ECM assembly and myotube formation *in vitro*.^59^ Perlecan whole-KO mice exhibit perturbed development of the skeletal system, and while they do initially form basement membranes, these are unstable and rapidly deteriorate due to shear stresses, particularly in the heart and vasculature.^53,60,61^ With the interpretation that Perlecan is a re- quiescence cue for MuSCs during regeneration, the predicted phenotype held true when a transient knockdown of the protein *in vivo* led to a surplus of myogenic cells going into 7 DPI and a reduction at the 14 DPI endpoint. Without Perlecan, transient amplifying myoblasts production was escalated at the expense of MuSC population maintenance. Strikingly, despite a temporary knockdown of Perlecan where the levels in the tissue eventually recovered (immunostaining not shown), the reduced MuSC pool at 14 DPI was also reduced in the proportions that re-quiescence and home to their niche, indicating that the timing of Perlecan presentation (between 4-7 DPI) is critical.

The Perlecan knockdown further presented a unique myofiber phenotype, a transient increase in smaller myofibers forcibly forming under the same basal lamina. We observed a self-correction by 14 DPI (speculatively through lateral fusion), yet there was a skew towards type IIB at the expense of type IIA myofibers. However, we did not note a hinderance in the myofiber maturation at 7 DPI. Thus, it is interesting to consider that the surplus of differentiating cells was somehow able to skew myonuclear fate, perhaps influenced by lateral fusion or the differing gene regulatory networks of division dependent versus independent fusion.^62^ Furthermore, knockdown of Perlecan led to an abnormal fibrotic response. While transient fibrosis represents an important component to regeneration^43^, the disruption of this ECM building block led to collagen I scarring at 7 DPI, that was corrected by 14 DPI (data not shown). As previously mentioned, the basal lamina represents a physical barrier (occasionally termed protective) to other mononuclear cell types with respect to the myofiber and MuSCs, which even during regeneration are maintained in their sheaths (i.e., “ghost fibers”).^20,21,63^ The weakening of this barrier through si*Hspg2* may have triggered a compensatory response similar to the observed fibrosis in aged skeletal muscle.^64–67^ As well, this mirrors the tissue instability brought on in Duchenne muscular dystrophy which sees elevated ECM proteins including Perlecan.^68^

Circling back to Perlecan signaling, the corresponding receptor (according to the database used) is α-dystroglycan (*Dag1*) where the binding is tight (dependent on glycosylation and calcium), acting through the laminin-globular domains of Perlecan.^69–73^ Moreover, literature indicates a binding to the integrin β1 subunit, also present on the MuSC surface.^72,74,75^ Both binding sites are found on the domain V of Perlecan, which can be produced recombinantly as a bioactive ligand and was named Endorepellin for its anti-angiogenic effects.^76^ Endorepellin was able to prevent differentiation in mini-IDLE and subdue the MuSC population at 7 DPI in the knockdowns, demonstrating it as a genuine MuSC ligand. Admittedly, effects were observed at high ligand concentrations, which may suggest an advantage to full-length Perlecan; but it can be challenging to study due to its dynamic/context dependent nature. ^58^

The Perlecan myofiber-MuSC signaling axis shows to be dysregulated in the aged skeletal muscle, with again many aspects of this phenotype being similar to the si*Hspg2* treatment, and as such contributes to our knowledge of the aging ECM. While the differential expression appears minimal, it is increased from myocytes/myonuclei suggesting that not only is synthesis lowered but also shorter lasting which leads to the pronounced difference at the protein level. We thus propose that reduced Perlecan signaling in aged muscle at the critical point of late-stage regeneration lends to poor MuSC re-quiescence and contributes to population decline over time and multiple bouts of regeneration. Curiously, a different study reported a rise in Perlecan levels with age in the mouse gastrocnemius and a drop-off after 22-months using an immunolabelling method, concurrent with increased apoptosis of capillary cells.^77^ This may reflect a difference in age/methods used or instead how Perlecan is dysregulated/used to compensate in different compartments of the muscle ECM. Nonetheless, supplementation of this signaling with Endorepellin was sufficient to rescue the aged MuSC population at 14 DPI and promote the quiescence state, offering a new therapeutic strategy for modifying MuSC behavior in aged muscle (which already has a track-record as a systemically administered treatment for stroke and Alzheimers with its additional neuroprotective properties^78,79^).

## Materials and Methods

### Animal use protocols and ethics

For experiments involving mini-IDLE, the *in vitro* recombinant mouse endopellin studies, and *in vivo* perlecan protein assessment, animal use protocols were reviewed and approved by the local Animal Care Committee (ACC) within the Division of Comparative Medicine (DCM) at the University of Toronto. All methods in this study were conducted as described in the approved animal use protocol (#20012838) and more broadly in accordance with the guidelines and regulations of the DCM ACC and the Canadian Council on Animal Care. 129-Tg(CAG-EYFP)7AC5Nagy/J (Actin-eYFP) mice^80^ were purchased from the Jackson Laboratory by the lab of Dr. Derek van der Kooy and a breeding pair was shared with our group to start a colony (6- week old females were paired with 8-week old males and their cages). Gt(ROSA)26Sortm1.1(CAG-cas9*,-EGFP)Fezh/J^81^ mice were purchased from the Jackson Laboratory by the lab of Dr. Jason Moffat and transferred to our group for hindlimb tissue collection and myoblast line derivation. For these studies, young male mice were between 4-5 months and aged mice between 24-26 months old.

Animals were handled according to the guidelines of the European Community, adhering to the principles of the 3Rs (Replacement, Reduction, Refinement). Protocols were approved by the ethics committee of the French Ministry, under the reference number APAFIS #46396- 2023111418593412 v4. Mice were housed under specific pathogen-free conditions in individually ventilated cages, with a 12-hour light/12-hour dark cycle, and were provided with ad libitum access to water and food. All *in vivo* experiments were conducted using 12-week-old or 24-month-old male C57BL/6J mice obtained from Charles River Laboratories.

### Murine muscle stem cell magnetic-activated cell sorting (MACS) and primary myoblast derivation

MuSCs were prospectively isolated from mouse hindlimb muscle using a MACS-based method as previously described by our group.^29,82^ Briefly, ≈1 gram of minced hindlimb muscle (with scissors on ice) was placed into a 15mL tube and digested for 1-hour at 37 °C on a rocker with 630 U/mL Type 1A collagenase from clostridium histolyticum in 7 mL of high-glucose DMEM. Afterwards, 440 U of Type 1A collagenase was added along with Dispase II and DNAse I (100 µg/mL) for another hour at 37 °C. The sample was further processed by slowly passing through a 20 G needle 15 times and then resuspending in 7 mL of FACS buffer (**Table S3**). The sample was then filtered through a 70 µm cell strainer followed by a 40 µm cell strainer. The sample was centrifuged at 400 g for 15 minutes and the pellet was resuspended in 1 mL of 1X red blood cell (RBC) lysis buffer for 8 minutes (**Table S3**) and then washed with FACS buffer. The cells were then incubated in a 4 °C fridge for 15 minutes in 100 µL of MACS buffer (**Table S3**) and 25 µL of lineage depletion microbeads from the MACS Satellite Cell Isolation Kit. The sample was then depleted of the lineage positive cells by flowing the solution through a MACS LS column in a magnetic field. This depletion was repeated once more with a second new column and another 25 µL of lineage depletion microbeads in 100 µL of buffer (with incubation again). A third and final new column was then used for a positive selection with anti-integrin α-7 microbeads (25 µL with 100 µL of MACS buffer and incubation) where here the MuSC are bound inside the column and must be removed from the magnetic field and flushed with 4 mL of buffer over an empty tube. After flushing the cells out of the column and spinning down the pellet (400 g x 5 min), the enriched integrin α-7^+^ MuSCs were either seeded to myotube templates (below) or plated *in vitro* to establish primary myoblast lines. In the case of the latter, cells were placed in collagen I coated dishes (1:8) with SAT10 media (**Table S3**). A full media change was performed 48 hours later, with half media changes every 2 days thereafter. Cells were grown to 70 % confluency and passaged a minimum 5 times to establish a primary mouse myoblast line used for experiments. Cells were maintained at 37 °C and 5 % CO2.

### mini-IDLE culture assay

Generation of murine myotube templates and subsequent seeding of fresh adult MACS-enriched MuSCs has been described in detail in our original publication.^29^ Briefly, pluronic acid-coated (Sigma-Aldrich, #P2443) microwells were dried and then 5 mm discs of Cellulose paper (MiniMinit) were added. A 0.8 U/mL thrombin solution (in PBS) was diffused into the paper discs and left to dry at RT. Meanwhile, primary myoblasts were mixed into an ECM-slurry containing 40 % DMEM, 40 % Fibrinogen (10 mg/mL in 0.9 % wt/vol NaCl) and 20 % Geltrex^TM^ at a concentration of 25,000 cells per 4 µL. The mixture was then diffused into the paper discs and left to gel at 37 °C for 5 minutes. 200 µL growth media (GM, **Table S3**) was added for the first 2 days (Day -2 to 0). On Day 0 of differentiation, the GM was fully removed and exchanged to differentiation media (DM, **Table S3**). Half media changes with DM were performed every other day moving forward. For the co-culturing experiment, ECM-only tissues were generated with a similar protocol without myoblasts added to the ECM-slurry. These tissues were then placed in a 48-well microwell, together with a Day 5 myotube template, in a 300 µL total volume of DM.

On Day 5 of myotube template differentiation, tissues were removed from the 96-well plate using tweezers and placed in an ethanol-sterilized plastic container containing long strips of polydimethylsiloxane (PDMS) sitting on top of a moist paper towel. Quickly, 4 µL of the MuSC solution (SAT10 media replete of FGF2) containing ≈ 500 MuSCs was placed onto each myotube template and gently spread over the tissue surface using a cell-spreader. The plastic container with myotube templates seeded with MuSCs was then sealed and placed in the 37 °C incubator for 1 hour. Tissues were then returned to their individual wells containing DM.

### Mass spectrometry

#### Preparation of tissue lysates

Myotube templates or ECM-only tissues were cultured to Day 5. The media was then removed and the tissues were washed quickly, three times, with warm PBS. The tissues were then digested for 90 minutes in 150 µL of PBS with 15 µg of Trypsin. The lysate was then spun down twice for 6 minutes at 500g followed by a filtering step using a 40 µm mesh to remove cells and large debris. The resulting samples were then reduced by incubation in 10 mM tris(2-carboxyethyl)phosphine (TCEP) (final concentration) at 70 °C for 30 minutes, and then alkylated with 20 mM iodoacetamide (IAA) (final concentration) in the dark at room temperature for 30 minutes. Afterwards, trypsin (1:40) was added and the samples were incubated at 37 °C for 90 minutes. Trypsin (1: 40) was added again and the samples were incubated at 37°C for 16 h. The digests were then quenched by the addition of 1 μL of concentrated formic acid.

#### Preparation of conditioned media

Myotube templates or ECM-only tissues were cultured to Day 5. The media was then removed and the tissues were washed quickly, three times, with warm PBS. PBS was completely removed and replaced by 120 µL of high-glucose DMEM with 4% ACA. 24 hrs later the conditioned media was collected and centrifuged at 14,000 g and washed three times with 200 μL ammonium bicarbonate using Amicon™ Ultra 0.5 mL centrifugal filter devices (Millipore, #Z677094-24EA) with a cut-off mass of 3 kDa. The resulting protein sample was resuspended in 55 μL of 0.1 M ammonium bicarbonate. Proteins were then reduced in 10 mM TCEP (final concentration) at 70 °C for 30 minutes, alkylated with 20 mM IAA (final concentration) in the dark at room temperature for 30 minutes. Afterwards, trypsin (1:40) was added and incubated at 37°C for 90 minutes. Trypsin (1: 40) was added again followed by incubation at 37°C for 16 h. The protein digests were then quenched by the addition of 1 μL of concentrated formic acid.

#### NanoRPLC-ESI-MS/MS Methods

Nanoflow reversed-phase liquid chromatography (NanoRPLC) was performed on an EASY-nLC 1200 ultra-high-pressure system coupled to a Q-Exactive HF-X mass spectrometer equipped with a nano-electrospray ion source (Thermo Fisher Scientific). Peptide samples were automatically loaded onto a C18 trap column (100 μm i.d. × 5 cm) and separated by a C18 capillary column (75 μm i.d. × 15 cm). The trap column and the analytical column were both packed in-house with 1.9 μm, 120 Å ReproSil-Pur C18 reversed phase particles (Dr. Maisch GmbH). Two mobile phases (A: 0.1 % (v/v) FA and B: 80/20/0.1 % ACN/water/formic acid (v/v/v) (Acetonitrile, ACN, Fisher Chemicals, A9551)) were used to generate a 130 minute gradient with a flow rate of 250 nL/minutes (kept at 3 % B for 5 minutes, 90 minutes from 3 % to 30 % B, 20 minutes from 30 % to 45 % B, 1 minute from 45 % to 95 % B and 14 minutes to kept at 95 % B). MS was operated in full scan (120,000 FWHM, 375-1575 m/z) and MS/MS scan (60,000 FWHM, 200-2000 m/z) with HCD collision energy of 27 %. The maximum IT for MS and MS/MS was set as 50 and 250 ms, respectively. The electro-spray voltage was 2.0 kV, and the heated capillary temperature was 275 °C. All the mass spectra were recorded with Xcalibur software (Thermo Fisher Scientific). Lysine-C was purchased from New England BioLabs.

#### Database Search and Quantification Analysis

Maxquant 2.0.3.1 were applied for database searching against the Uniprot_mus_musculus database (updated on 20/07/2022, 21989 entries). The parameters of database searching included: up to two missed cleavage were allowed for full tryptic digestion; carbamidomethylation (C) as a fixed modification and oxidation(M), acetyl (protein N-term) as variable modifications; label free quantification (LFQ) and match between runs (MBR) were activated. Peptide spectral matches (PSM) were validated based on q-value at a 1 % false discovery rate (FDR). Default settings were used for other parameters. Quantification was conducted by Perseus 1.6.15.0. Within Perseus, the initial proteomic dataset was filtered removing potential contaminants and reverse hits. Further filtering of the dataset was carried out considering for each protein a number of valid LFQ values equal to or higher than 3 in each group. LFQ intensities were log2-transformed, and missing values imputed using random numbers from an ideal gaussian distribution (width = 0.3; down shift = 1.8). Finally, the comparison between the myotube templates vs ECM-only conditions was carried out selecting significant proteins with values of S0 = 0.5 and Student’s t-test with FDR = 0.01.

### Single-nuclei RNAseq

Using the protocol from Dos Santos et al. (2021)^83^, we isolated tibialis anterior (TA) muscles from young and aged mice at different timepoints after a cardiotoxin-induced injury (4 and 7 DPI). After dissection and chopping of the tissue into 2-3 mm sized pieces, the muscle was transferred to a lysis buffer (substituting Nonidet^TM^ P40 for IGEPAL® CA-630 from the original protocol) (**Table S3**) and homogenized using a 5 mL Douncer. The lysate was filtered through 70 and 40 µm strainers, centrifuged, and resuspended in wash buffer (**Table S3**) with DAPI stain, and then DAPI+ nuclei were sorted using FACS. The number of nuclei was determined with a Luna FL counter (Logosbio) to obtain an expected cell recovery population of 8,000 cells per channel, loaded on a 10x G chip and run on the Chromium iXsystem (10x Genomics) according to manufacturer’s instructions. Single-cell RNA-seq libraries were generated by the Cancer Genomic Platform (CRCL, Lyon) with the Chromium Single Cell 3′ v.3.1 kit(10x Genomics, no. PN- 1000121) and sequenced on the NovaSeq 6000 platform (Illumina) to obtain around 50,000 reads per cell.

#### Data analysis

The FASTQ files and gene-barcode matrix were generated using cellranger software (v7.0.0) and the RNA STARSolo tool from Galaxy .^84^ The following parameters were used:

--genomeFastaFiles Mus_musculus.GRCm39.dna.primary_assembly.fa

--genomeSAindexNbases 14

--sjdbGTFfile Mus_musculus.GRCm39.108.gtf

--sjdbOverhang 100

--soloCBwhitelist 3M-february-2018.txt

--soloUMIdedup 1MM_CR

--soloCBmatchWLtype 1MM_multi_pseudocounts

--soloStrand Forward

--soloFeatures Full

--soloUMIfiltering MultiGeneUMI_CR

--soloCellFilter EmptyDrops_CR 15000 0.99 10.0 45000 90000 500 0.01 20000 0.01 10000

--soloOutFormatFeaturesGeneField3 “Gene Expression”

Matrices were then loaded into RStudio and underwent standard quality control using Seurat^85^, and doublet removal with DoubletFinder^86^. After clustering and marker identification, cells were manually annotated using a combination of the differentially expressed markers and the following:

MuScs: Pax7

Activated MuSCs: Pax7, Lockd

Myocytes/fusing: Myog, Jam3

Immunomyoblasts: Pax7, Ncam1, Ctsb, Ctss

FAPs: Pdgfra

Rian+ nuclei: Rian

Nascent myonuclei: Myh3, Myh8

Type IIX: Myh1

Type IIB: Myh4

Type IIA: Myh2

NMJ: Etv5, Ache, Chrne

MTJ: Col22a1

ECs: Pecam1, Cdh5 Macrophages: Ctsa, Ctss Dendritic: Flt3

T and B cells and NK cells: Skap1, Cd8a, Cd11b

SMMCs: Myh11, Acta2

For data integration, individual Seurat objects (individual DPI/Age) were imported into Python and assembled with scANVI^87^ using the top 5000 genes with DPI and Age as categorical covariate keys. The integrated dataset was furthermore exported for continued analysis in RStudio with Seurat. Lastly, the CellChat tool was used for CCCI analysis.^34^ CellChat objects were created from Young and Aged subsets of the scANVI-integrated Seurat object, and the CellChatDB.mouse library of receptor-ligand interactions. Communication probabilities were computed from over expressed interactions and the population size parameter set to TRUE.

### Western blots

#### Protein lysate preparation

To identify and quantify specific proteins in TA muscles or myotube templates, we conducted western blot analysis. First, a lysate buffer (**Table S3**) was mixed with dissected TAs at 100 mg/mL or 200 µL per 25 myotube templates. The sample was disrupted vigorously with a motorized pestle gun and then incubated on ice for 30 minutes followed by a 10 minute x 10,000 g centrifugation at 4 °C. Protein concentration for each sample was then determined using a BCA assay (Thermo- Fisher, #23227).

#### SDS-PAGE Gel

Components for the 8 % resolving gel (**Table S3**) were mixed and pipetted into an empty gel cassette, carefully overlaid with isopropanol to cover the top, and then allowed to polymerize for 30 minutes. The isopropanol as removed and the components for the stacking gel (**Table S3**) were mixed and pipetted on top of the polymerized resolving gel. The comb was inserted, and the stacking gel was allowed to polymerize for 10 minutes. The cassette containing the gel was then wrapped in a moist paper towel and stored overnight at 4 °C. The next day, the gel was loaded into the buffer tank with running buffer (**Table S3**). The wells were loaded with the HiMark™ Pre- stained Protein Standard ladder or with protein samples (40 µg for TA lysate or 20 µg for myotube template lysate) that were diluted in 50 mM Tris pH 8.0 buffer and 6X SDS sample buffer. The gel was run at 80V for 3 hours. Afterwards, the gel was transferred to a nitrocellulose membrane with wet a transfer (see buffer in **Table S3**) at 60V for 1 hour. The membrane was then stained and imaged with Ponceau solution (Bio-Rad ChemiDoc^TM^ Imaging System), washed, blocked with 5 % skimmed milk (SM) in TBST buffer (**Table S3**) for 45 minutes, and then incubated overnight with primary antibodies (**Table S2**) in 5% SM-TBST in a 4 °C fridge. The membranes were then washed 3 x 10 minutes with TBST, incubated for 1 hour with HRP-conjugated secondary antibodies (**Table S2**) in 1% SM-TBST, and then chemiluminescence signals were captured also using a Bio-Rad ChemiDoc^TM^ Imaging System after treatment with the SuperSignal™ West Dura Extended Duration Substrate.

### Lentiviral sgRNA production

#### sgRNA design and plasmid assembly

The pLCKO backbone plasmid bacterial agar stab was purchased from Addgene (#73311), streaked on LB agar plates with ampicillin, and a single colony inoculated in liquid culture overnight at 37 °C. Plasmid DNA was then purified using the QIAprep Spin Miniprep Kit (Qiagen, #27104). 20 bp sgRNA sequences were designed using CHOPCHOP v3^88^ (**Table S4**) and ordered as complimentary oligos (Integrated DNA Technologies, Custom Oligos). Afterwards, the pLCKO backbone was digested using BfuAI and NsiI overnight at 37 °C followed by calf intestinal alkaline phosphatase (CIP) treatment for 2 hours at 37 °C. The reaction was run on a 1 % agarose gel at 100 V for 90 minutes. The 7500 bp vector was cut out and purified using the QIAquick Gel Extraction Kit (Qiagen, #28704). Forward and reverse oligos for each sgRNA were annealed together with T4 Polynucleotide Kinase in a thermocycler with the following conditions: 37 °C for 30 minutes, 95 °C for 5 minutes and ramp down to 25 °C at 5 °C/minute. Annealed oligos were diluted 1:250 and ligated to the pLCKO vector using T4 ligase overnight at 16 °C. The reaction then underwent standard heat shock transformation using chemically competent C3040 bacteria and incubated overnight on LB agar plates with ampicillin. A single colony was then expanded in liquid culture, the plasmid DNA purified.

#### Lentiviral packaging

For each sgRNA, 5 x 10^6^ 293T cells were plated in 10 cm plates in low-antibiotic growth media (**Table S3**). The cells were transfected 24 hrs later. A mixture of the ligated vector (6 µg), the psPax2 packaging plasmid (6 µg, Addgene, #12260), the pMD2.G packaging plasmid (2 µg, Addgene, #12259), and 3X polyethylenimine was added dropwise to the culture and incubated for 18 hours at 37 °C. Next, the plate was washed with warm PBS and replaced with 10 mL of low-antibiotic growth media. After another 24 hours, the viron-containing media was collected, centrifuged at 300 g x 10 min and passed through a 0.45 µm filter to remove any cells prior to aliquotting into polypropylene tubes, which were stored at -80 °C until use.

### T7 endonuclease assay

A T7 assay was performed to verify Cas9 activity at the appropriate target site for each of the sgRNAs. Cas9 expressing myoblasts were seeded into a 6-well plate at 3 x 10^5^ cells / well. For editing in myoblasts, cells were left to equilibrate for 48 hours prior to incubating with lentiviral sgRNAs for 18 hours (viral load was proportional to the medium volume/cell ratio in mini-IDLE). For editing in myotubes, after 48 hours in growth media, the cultures were exchanged to DM for 72 hours. The cells were then transduced with lentiviral sgRNAs for 18 hours. 24 hours after virus removal, the myoblast and myotube cultures were collected and the genomic DNA extracted using the DNeasy Blood & Tissue Kit (Qiagen, #69504). Primers to amplify 500-1000 bp products (**Table S5**) were designed for each sgRNA/target site and the editing efficiency assessed in a semi-quantitative manner using the EnGen® Mutation Detection Kit (New England Biolabs, #E3321S).

### Myotube template transduction

For ligand gene editing on Day -1, 24 hours after myoblast seeding the GM was completely removed and replaced with 170 µL of GM with 10 µL (Low), 30 µL (Medium) or 50 µL (High) of the lentiviral sgRNA stock solution. After 16-18 hours post-infection, tissues were quickly washed three times with GM and then incubated in GM for another 8 hours before switching to DM. When editing on Day 4, a full media change was done to replace the GM with 170 µL of DM with 10 µL (Low), 30 µL (Medium) or 50 µL (High) of the lentiviral sgRNA stock solution. 16- 18 hours post-infection, tissues were quickly washed three times with DM and then incubated in DM for another 12-16 hours before seeding the MuSCs.

### MTS assay

The metabolic activity of myotube templates was quantified by adding 200 µL of fresh DM with 20 µL of the MTS tetrazolium compound (Abcam, #ab197010) for 2 hours at 37 °C with mixing every 30 minutes. The entire solution was then pipetted into a clear 96-well plate and the OD at 490 nm quantified with a spectrophotometer (Tecan, Infinite M200 Pro). The “media + MTS” negative control was subtracted as background from all OD values for each repeated experiment.

### *In vitro* sample fixation and immunolabelling

As previously described^29^, media was removed from myotube template or 2D culture samples followed by 3x quick washes with PBS followed by fixation with 4 % paraformaldehyde (PFA) for 12 minutes at RT. Fixative was removed followed by 3 x 10 minute washes with PBS and then samples were blocked and permeabilized (**Table S3**) for 30 minutes at RT. Afterwards, primary antibodies (**Table S2)** diluted in blocking solution were added to samples, which were then incubated overnight at 4 °C. After 3 x 10 minute washes with PBS, samples were incubated for 45 minutes at RT with secondary antibodies diluted in blocking solution (**Table S2**), followed by 3 x 10 minute washes with PBS. For EdU studies, labelling was conducted using the Click-iT™ Plus EdU Alexa Fluor™ 555 Imaging Kit (Invitrogen, #C10638). EdU was added directly to the culture media (10 µM) and was refreshed every 24 hours until the indicated endpoint.

### *Hspg2* siRNA and Endorepellin *in vivo* studies

Mice were anesthetized using isoflurane inhalation in preparation for muscle injury. Acute muscle injury was induced by intramuscular injections of 50 μL of cardiotoxin (CTX; 10 μM)^89^ into the Tibialis Anterior (TA) muscle. All in vivo experiments were performed across two independent experiments, each consisting of N=3 mice. TriFECTa® RNAi Kits with siCtl and si*Hspg2* were obtained from Integrated DNA Technologies (IDT). A total of 0.8 nmol of siCtl or si*Hspg2* in a 60 μL volume, was injected intramuscularly into each TA muscle at two time points: 2 days and 3 days after CTX injury. For each siRNA injection, a RNAiMax mix containing Opti-MEM and RNAiMax was prepared in one tube, and a second mix containing siRNA and Opti-MEM was prepared in a second tube. Therefore, the first mix was added to the second one and gently homogenized.

#### In vivo Endorepellin experiment

For each TA muscle, 6 μg of Endorepellin diluted in 60 μL volume of PBS was injected using intra-muscular injections. The control conditions were injected with PBS only.

### RNA extraction, cDNA synthesis and RT-qPCR

A minimum of 2 x 10^5^ cells were collected per sample for RNA extraction using the Nucleospin RNA II kit following the manufacturer’s protocol. RNA concentration was assessed using a Nanodrop spectrophotometer. Complementary DNA (cDNA) was synthesized using the High-Capacity Reverse Transcription Kit (Thermo Fisher Scientific, #4368814). The cDNA was then utilized for quantitative PCR (qPCR) performed with SYBR Green Master Mix and run on a LightCycler 480 for 40 cycles. Primer sequences are provided in **Table S6**. All samples were duplicated, and transcript levels were normalized to a housekeeping gene for relative abundance. The relative mRNA levels were calculated using the 2^−ΔΔCt method. The ΔCt values were obtained from Ct values normalized to the housekeeping genes Gapdh or Hprt levels in each sample. Ct values between 18 and 28 were considered for analysis.

### Isolation of TA muscle and histological staining

Mice were euthanized through isoflurane asphyxiation followed by cervical dislocation. TA muscles were harvested, weighed, submerged in OCT, and then frozen in isopentane cooled in liquid nitrogen. Frozen muscles were stored at -80 °C. Muscle cryosections of 10 μm thickness were collected using an NX50 cryostat. Fixation of muscle sections was performed with 4 % PFA in PBS for 10 minutes at room temperature. Permeabilization was achieved using 0.1 M glycine and 0.1 % Triton X-100 diluted in PBS for 10 minutes at room temperature. After three washes in PBS, muscle sections were incubated with blocking buffer (3% BSA, 0.1% Triton X-100 in PBS) supplemented with M.O.M. Blocking reagent for 1 hour. Primary antibodies diluted in blocking buffer were then incubated overnight at 4 °C. Detection of primary antibodies was carried out using Alexa Fluor secondary antibodies diluted at 1:1000 in blocking buffer after three washes in PBS. Nuclei were counterstained with Hoechst at 1 μg/mL in PBS before mounting with fluorescent mounting medium.

### Image acquisition

For mini-IDLE images, as previously described^29^, confocal imaging was performed using the Perkin-Elmer Operetta CLS High-Content Analysis System and the associated Harmony® software. Data was collected using the 10x air objective (Two Peak autofocus, NA 1.0 and Binning of 1), and the 20x and 40x water immersion objectives (Two Peak autofocus, NA 1.0 and 1.1, and Binning of 1). All images were exported off the Harmony® software in their raw form. Later processing was performed using the ImageJ-BIOP Operetta Import Plugin available on c4science.^90^ In some cases, representative images were gathered with an Olympus FV-1000 confocal microscope and Olympus FluoView V4.2b imaging software with a 40x silicone immersion objective (NA 1.25; Olympus, #UPLSAPO40XS). For imaging of muscle cross-sections, entire muscle sections were assembled using a slide scanner ZEISS AxioScan7 using the 20x objectives. Muscle cross section images were processed and analyzed with ZEISS ZEN and FIJI software.

### Phenotypic data analysis

#### mini-IDLE MuSC analysis

For mini-IDLE phenotypic data analysis, CellProfiler^TM^ software was utilized as previously described.^29^ CellProfiler^TM^ version 4.2.150 was downloaded from the source website (www.cellprofiler.org) and installed on a PC (Intel Core i9-11900 @ 2.5GHz, 64.0 GB RAM, and 64-bit Windows 11 operating system). 25 max projected images were taken per tissue at 20x magnification. The channels were split, mononucleated DAPI^+^YFP^+^Pax7^+^ objects were extracted using the IdentifyPrimaryObjects module, and then counted. Morphology measurements of the identified cellular objects were recorded using the MeasureObjectSizeShape module. To quantify the proportion of MyoD^+^ cells, this fourth channel was overlayed over the extracted DAPI+YFP+Pax7+ objects (i.e. donor MuSCs) performed in the prior step. For YFP coverage, the YFP channel was isolated from 20x images and calculated using the MeasureAreaOccupied function. An average was created for each tissue by combining calculation from all images per tissue.

#### Myotube template analysis

For SAA coverage analysis, stitched images were prepared and analyzed with a previously published ImageJ macro^91^. The SAA signal was pseudocoloured in red, the threshold set to 0-45 and the tissue outline was selected using the oval tool. Myotube diameter was determined by manually measuring 20-30 random myotubes across all images taken for a specific tissue and calculating an average. To count nuclei and calculate fusion index, 6 max projected images were taken per tissue at 20x magnification. The channels were split, the myotube and nuclei signals were individually identified and overlayed to calculate the percentage of nuclei in myotubes. Z-line fraction was determined by converting 40x magnification images into max intensity projections and splitting the channels (SAA and actin) into individual TIFF files and using zlineDetection on MATLAB (R2022b)^92^ The recommended settings were used with the following modifications:

Dot Product Threshold = 0.77

Actin Segmentation Grid Size = 150 pixels

Actin Segmentation Threshold = 0.87

Noise Removal Area = 16 pixels

Skeletonization Branche Size Removal = 5

Results for each image were then averaged per tissue.

#### Muscle cross-section analyses

MuSCs were manually enumerated by identifying cells whereby nuclear Pax7 immunostaining colocalized with Hoechst staining. Collagen I and F4/80 immunostainings were quantified (semi-automatically?) using FIJI thresholding, and then reporting the % total area between the myofibers that was positive for each signal. Central myonuclei and the number of myofibers were quantified manually with the assistance of the FIJI cell counter tool.

The myofiber periphery was visualized by laminin-α2 immunostaining to then conduct cross-sectional analysis using the Open-CSAM ImageJ macro. Entire sections were analyzed, resulting in approximately 1500 myofibers counted for each tissue section.

### Statistical analysis

Statistical analysis was performed using the GraphPad Prism software and tidyverse packages in R. *In vitro* experiments were performed with 3 technical tissue/microwell replicates per experimental group and repeated on 3 independent occasions (e.g., n=9 technical replicates across N=3 biological replicates). *In vivo* experiments were performed with tissue collected from independent mice (e.g., N=5 mice). Please refer to **Table S7** for a specific breakdown of replicates per experiment and for documentation of statistical tests. Significance was defined as p < 0.001 for mini-IDLE assay experiments and p ≤ 0.05 for all other experiments.

## Acknowledgements

This work was supported by a CIHR CGS-D scholarship (E.J.), a MITACS Globalink Research Award (E.J.), a Michael Smith Foreign Study Supplement (E.J.), and a 4th year of PhD studies fellowship from la Fondation pour la Recherche Médicale (P.G.). The research conducted in the Gilbert Lab was supported by funding from the Medicine by Design Canada First Research Excellence Fund (MbDC2-2019-02) and the Canadian Institutes of Health Research (PJT- 186224). P.M.G. holds the Canada Research Chair in Endogenous Repair. The research conducted in the Le Grand lab was supported by funding from the Inserm, the CNRS, the Agence National pour la Recherche (ANR-19-CE13-0016, ANR-17-CE12-0010- 02), the Association Française contre les Myopathies/AFM-Téléthon (MyoNeurAlp2 project) and the European Joint Program for Rare Diseases (EJPRD JTC2019, Myocity project). Mass spectrometry/proteomics research was supported by a Precision Medicine Initiative fellowship to Y.H. and a Canadian Foundation for Innovation/Province of Ontario grant (36661) to A.R.W. We thank the Cancer Genomic Platform from the CRCL, in Lyon, France, for all the single nuclei RNA-seq sequencing.

## Author contributions

P.M.G. and F.L.G. conceived the project. E.J. and P.G. designed and performed research, analyzed data, and prepared figures. P.M.G., E.J. and F.L.G supervised the research and contributed to fundraising. All authors contributed to data interpretation. C.D. contributed to library preparation and sequencing. Y.H. designed and implemented the mass spectrometry/proteomic data collection pipeline, with support from A.R.W. O.M. and J.R. contributed to *in vivo* experiments. S.F. contributed to bioinformatics analyses. M.P.D.A. and S.A. provided resources and guidance for gene editing experiments. E.J., P.G., P.M.G. and F.L.G. wrote the manuscript. All authors reviewed, edited, and approved the manuscript.

---

**Figure S1.**
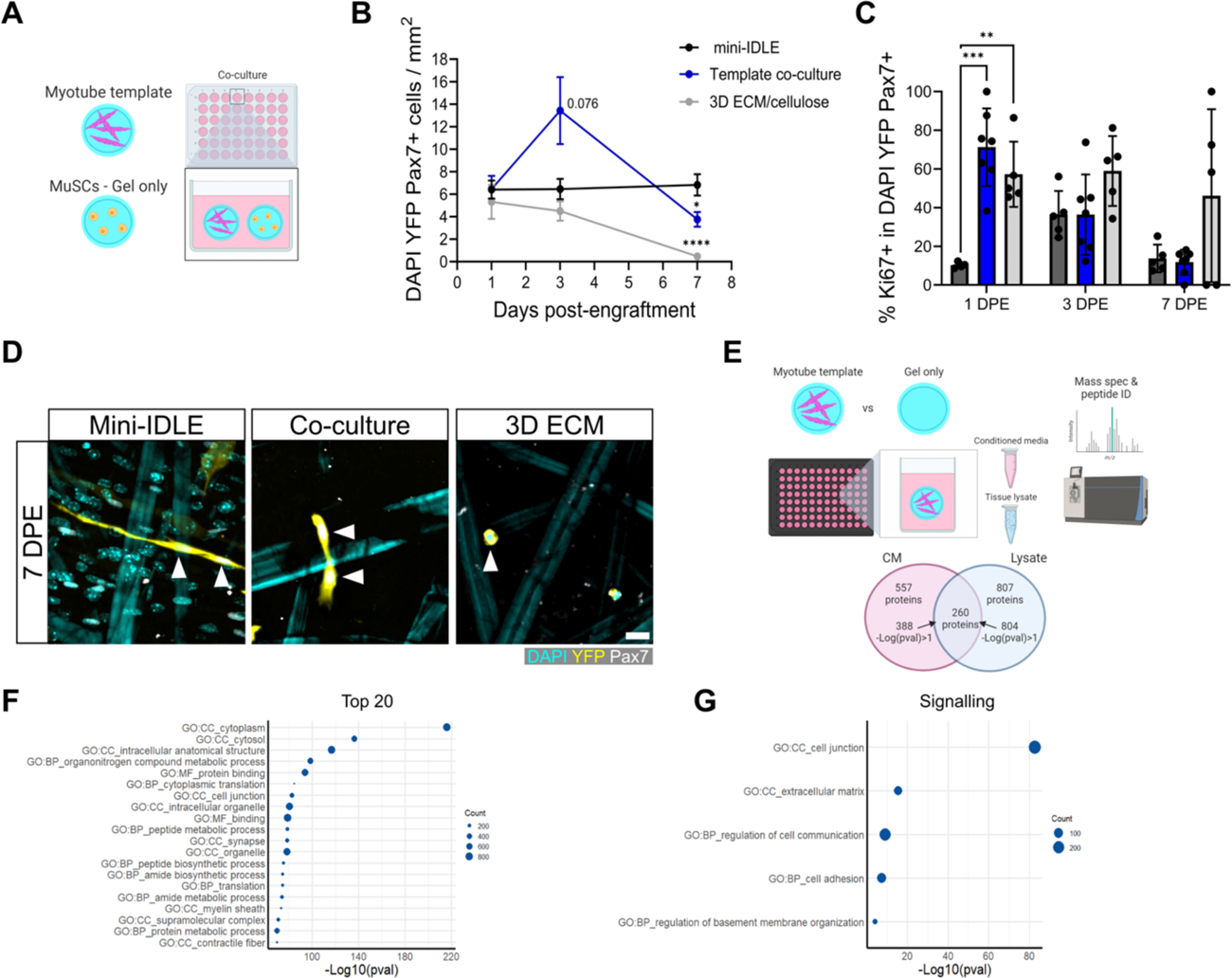
The mini-IDLE MuSC quiescence phenotype is irreplicable with diffused cues alone. **(A)** Schematic representation of the experimental setup (made with BioRender). **(B)** Quantification of mononuclear DAPI+YFP+Pax7+ cell density per mm^2^ at 1, 3, and 7 days post engraftment (DPE) across the mini-IDLE (black), template co-culture (blue), and 3D ECM-only (grey) conditions. n=4-7 tissues across N=2-3 independent biological replicates. Graph displays mean ± s.e.m.; one-way ANOVA with Dunnet’s test for each individual timepoint comparing against the mini-IDLE condition, *p=0.0117, ****p˂0.0001. **(C)** Bar plot showing the percentage Ki67+ in the DAPI+YFP+Pax7+ mononuclear cell population at each timepoint across the same conditions. n=4-7 tissues across N=2-3 independent biological replicates. Graph displays mean ± s.e.m. with individual technical replicates; one-way ANOVA with Dunnet’s test for each individual timepoint comparing against the mini-IDLE condition, **p=0.0018, ***p=0.0001. **(D)** Representative confocal images of donor cells in mini-IDLE, template co-culture, and ECM-only tissues at 7 DPE labelled for DAPI (cyan), YFP (yellow) and Pax7 (white, white arrows). Scale bar, 50µm. **(E)** Schematic of the experimental setup for mass spectrometry and a Venn diagram of the proteins identified in the conditioned media (CM) and myotube template lysate (made with BioRender). **(F-G)** Dot plots showing the top 20 GO terms **(F)** and 5 GO signaling terms **(G)** (in descending order) identified using all significantly identified proteins in the CM and lysate.

**Figure S2.**
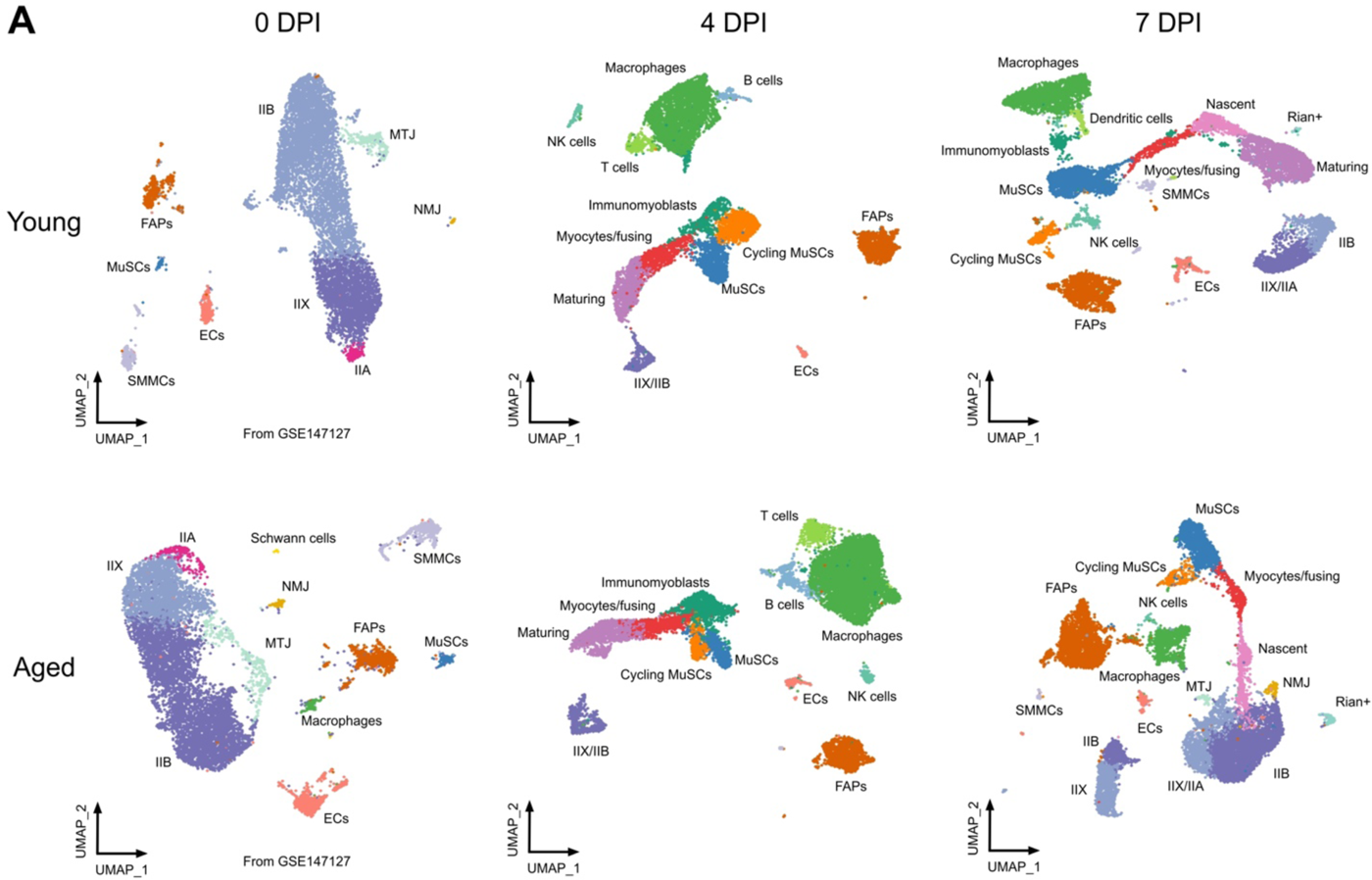
Clustering of individual snRNAseq datasets. Individual UMAP plots corresponding to 0, 4 and 7 days post injury (DPI) snRNAseq datasets collected from both Young and Aged with their respective annotated cell types. 0 DPI datasets from GSE147127.

**Figure S3.**
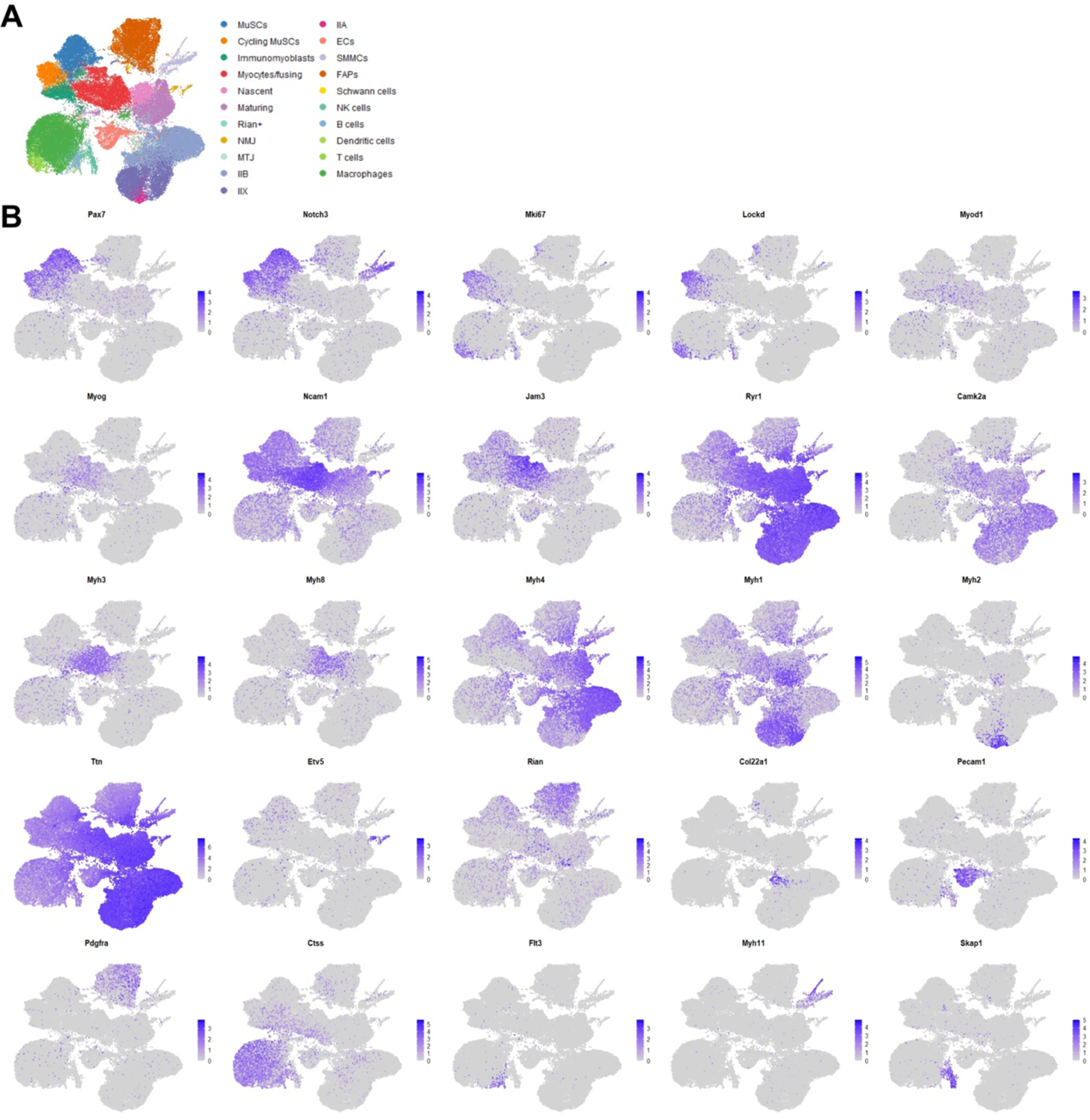
Gene expression in snRNAseq dataset of regenerating skeletal muscle. **(A)** Representative UMAP plot of the scANVI-integrated snRNAseq dataset with a color legend of the identified cell types. **(B)** Feature plots showing expression of selected genes involved in myogenesis, cell fusion, myonuclei specialization, and cell type identification.

**Figure S4.**
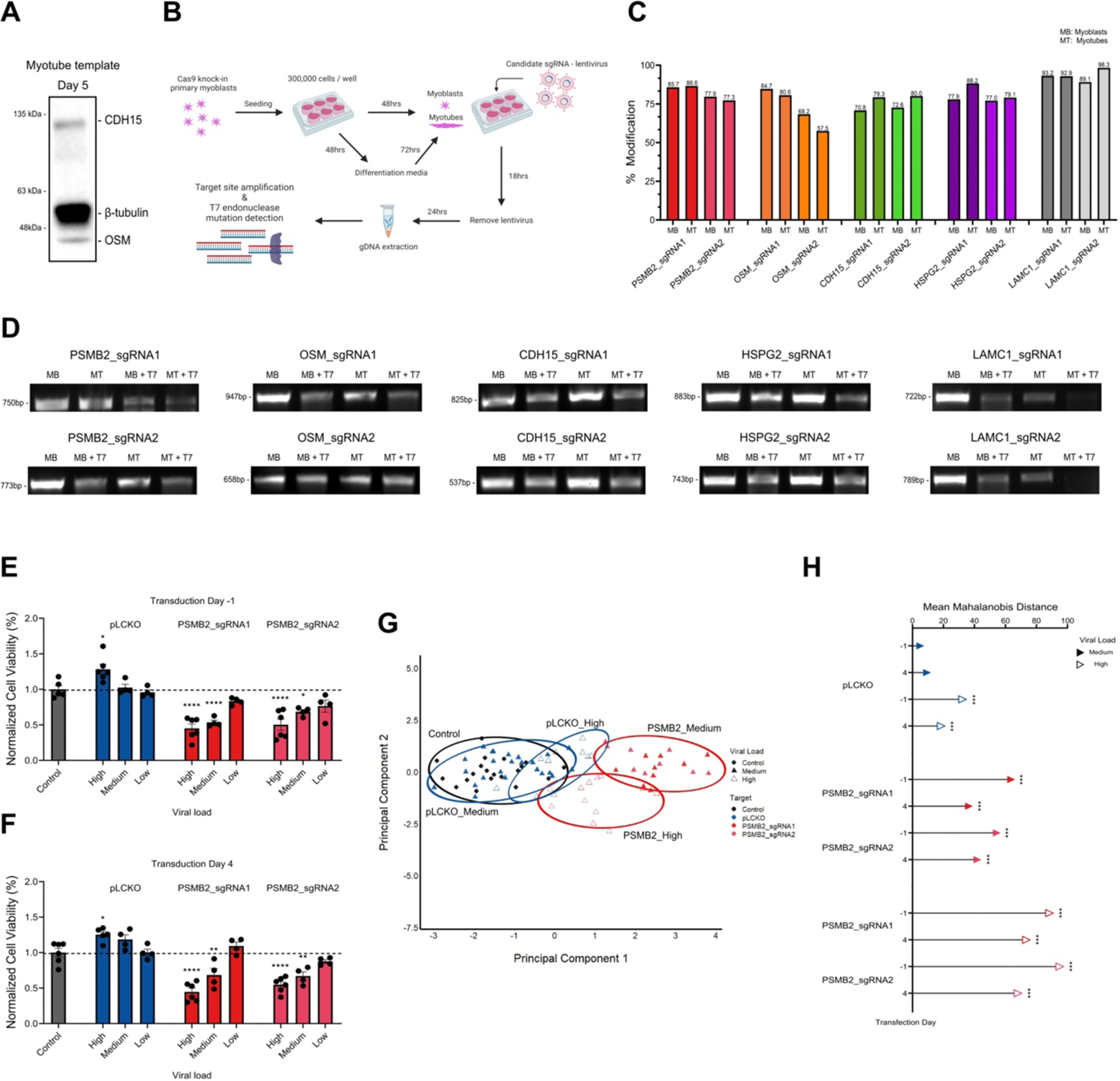
Validation of Cas9 editing and phenotype perturbation in mini-IDLE. **(A)** Representative chemiluminescent image of a western blot for Day 5 myotube template lysate confirming the presence of CDH15 (M- cadherin) and OSM (Oncostatin-M) proteins, along with β-tubulin. **(B)** Schematic representation of the experimental layout for testing Cas9 editing activity at sgRNA target sites in myoblasts and myotubes (made with BioRender). **(C)** Bar graph of the editing efficiency (% modification) of all sgRNAs tested in myoblasts (MB) and myotubes (MT). Concrete values, rounded to the first decimal point are indicated on the top of each bar. **(D)** Representative images of SYBR-safe stained PCR products of all sgRNA target sites without and with T7 endonuclease treatment (+T7), from myoblast and myotube samples. **(E-F)** Bar plots of normalized cell viability of myotubes templates on Day 5 of culture across different sgRNA plasmids delivered at low, medium and high viral loads on Day -1 (myoblasts) **(E)** or on Day 12 delivered on Day 4 (myotubes) **(F)**. Graphs display mean ± s.e.m. with individual technical replicates; one-way ANOVA with Dunnet’s test for each condition compared against control. n=4-6 tissues across N=2 independent biological replicates. *p=0.0110, 0.0106, ****p˂0.0001 (E); *p=0.0131, **p=0.0027, 0.0016, ****p˂0.0001 (F). **(G)** Principal component analysis plot showing the principal component 1 and 2 values for the average MuSC population features of every tissue analysed. Color and shape code for the target site/sgRNA and the viral load, along with lassos for each group that center around their respectively centroids. n=4-20 tissues across N=2-3 independent biological replicates. **(H)** Lollipop plot showing the mean Mahalanobis distance for each experimental group compared to the control centroid. n=4-20 tissues across N=2-3 independent biological replicates; p-values were calculated for Hotelling’s T-squared statistic, ***p˂0.001.

**Figure S5.**
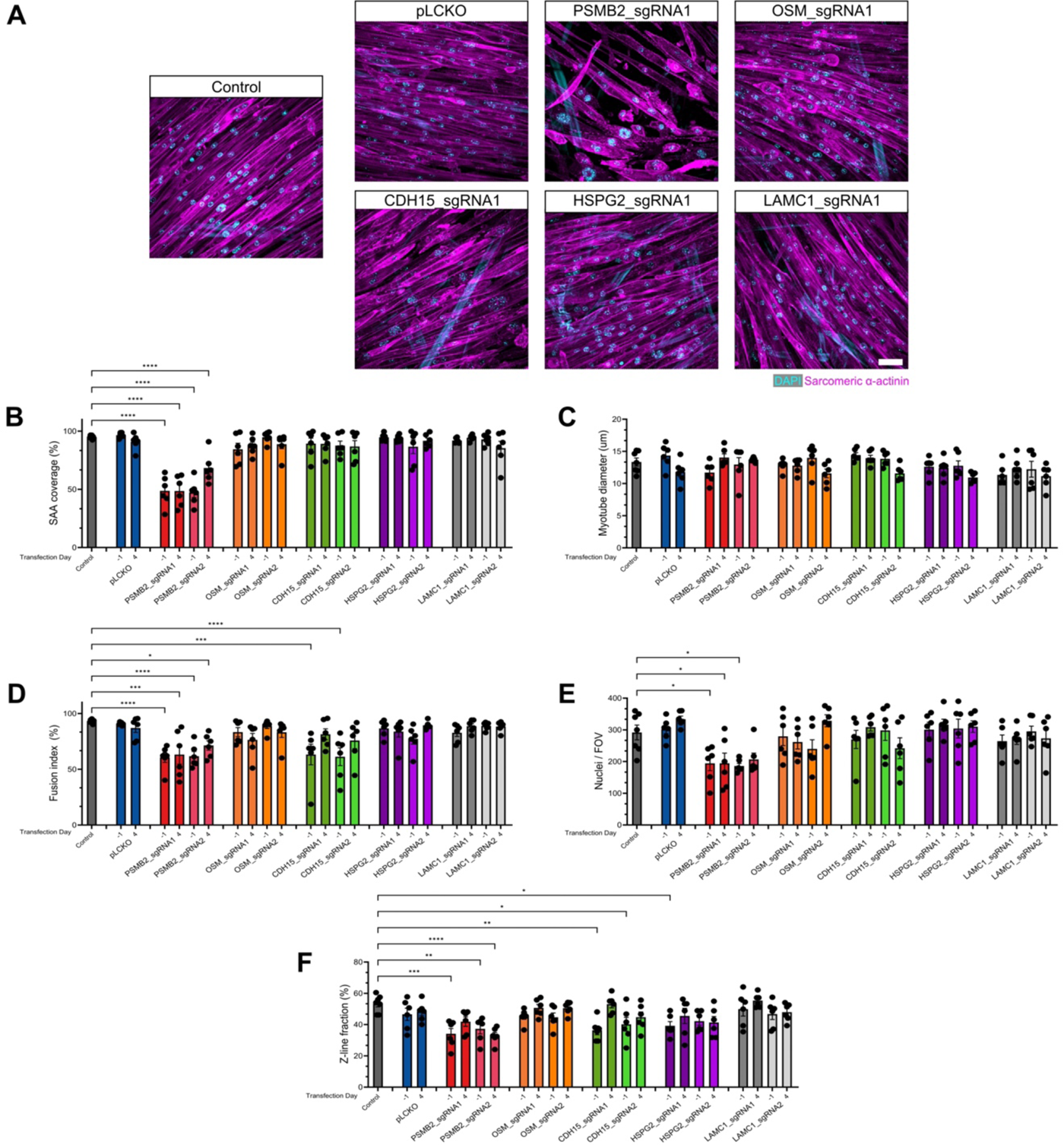
Myotube characterization under knock-down conditions. Representative confocal 40X images of myotubes on Day 12 after transduction on Day -1 with different sgRNAs; labelled with DAPI (cyan) and SAA (magenta). Scale bar, 50 µm. **(B-F)** Bar plot showing SAA coverage of myotube templates **(B)** average myotube diameters **(C)** percentage fusion index **(D)** average nuclei per field of view (FOV) **(E)** and percentage striated (z-line fraction) of myotubes **(F)** after transduction with different sgRNAs (color coded) on Day -1 or 4 (x-axis). Graphs display mean ± s.e.m. with individual technical replicates; one-way ANOVA with Dunnet’s test for each experimental condition compared against the control. n=6-7 tissues across N=2 independent biological replicates. ****p˂0.0001 (B); ns (C); *p=0.0140, ***p=0.0002, 0.0002, ****p˂0.0001 (D); *p=0.0386, 0.0389, 0.0177 (E); *p=0.0333, 0.0168, **p=0.0038, 0.0017, ***p=0.0002, ****p˂0.0001 (F).

**Figure S6.**
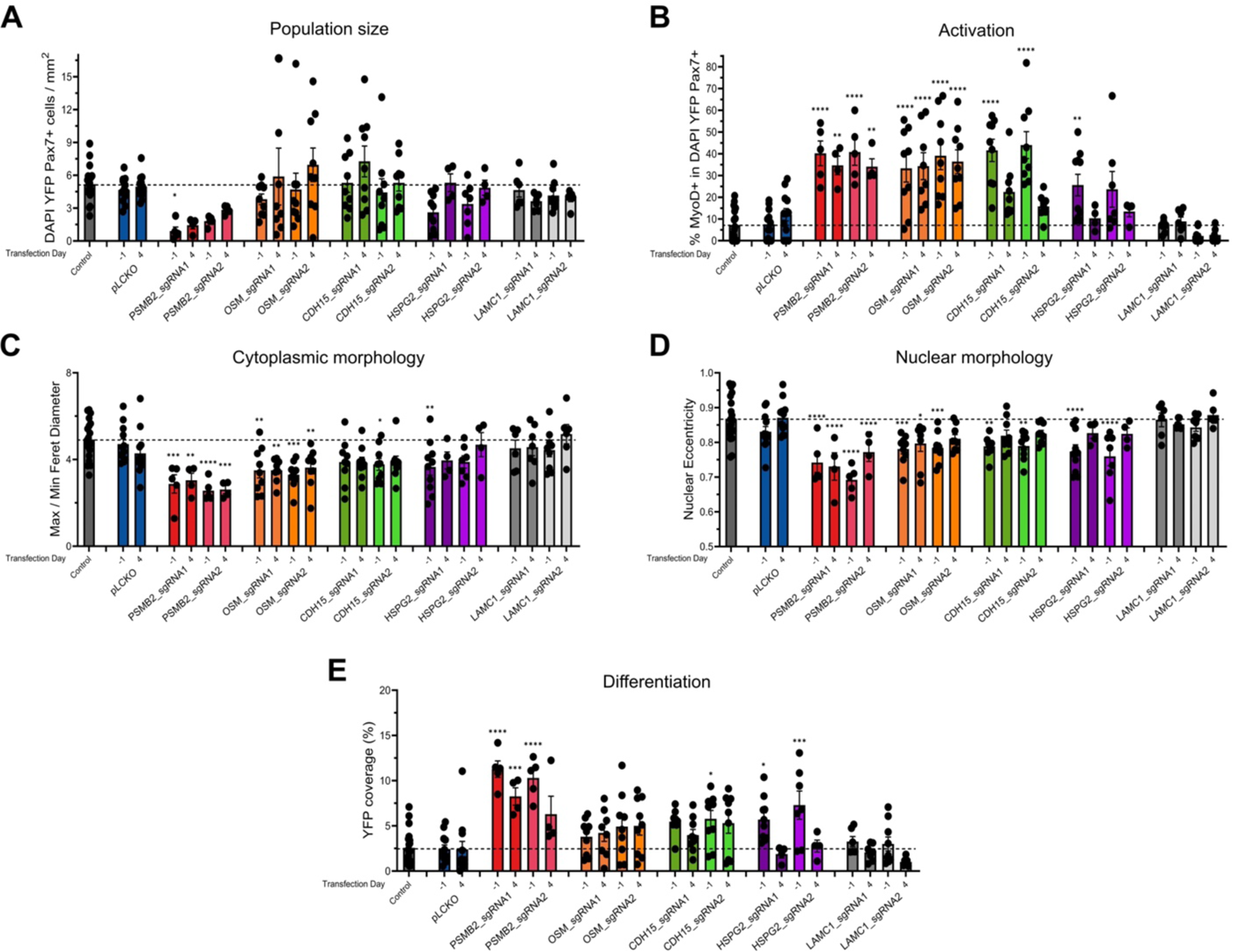
Breakdown of MuSC phenotype across candidate total knockdown conditions. **(A-E)** Bar plots showing mononuclear DAPI YFP Pax7+ cell density **(A)** percentage MyoD+ in the mononuclear DAPI YFP Pax7+ cell population **(B)** average max/min feret diameter of the mononuclear DAPI YFP Pax7+ cell population **(C)** average nuclear eccentricity of the mononuclear DAPI YFP Pax7+ cell population **(D)** and average YFP coverage per FOV **(E)** within tissue templates transduced with different sgRNAs (color coded) on Day -1 or 4 (x-axis). Graphs display mean ± s.e.m. with individual technical replicates and a dotted line to indicate the mean for control; one-way ANOVA with Dunnet’s test for each experimental condition compared against the control. n=4-20 tissues across N=2-5 independent biological replicates. *p=0.0446 (A); **p=0.0024, 0033, 0.0052, ****p˂0.0001 (B); *p=0.0321, **p=0.0036, 0023, 0.0043, 0.0084, 0.0072, ***p=0.0002, 0.0001, 0.0002, ****p˂0.0001 (C); , *p=0.0133, **p=0.0094, 0018, 0.0022, ***p=0.0004, 0.0007, ****p˂0.0001 (D); *p=0.0253, 0.0232, ***p=0.0007, 0.0004, ****p˂0.0001 (E).

**Figure S7.**
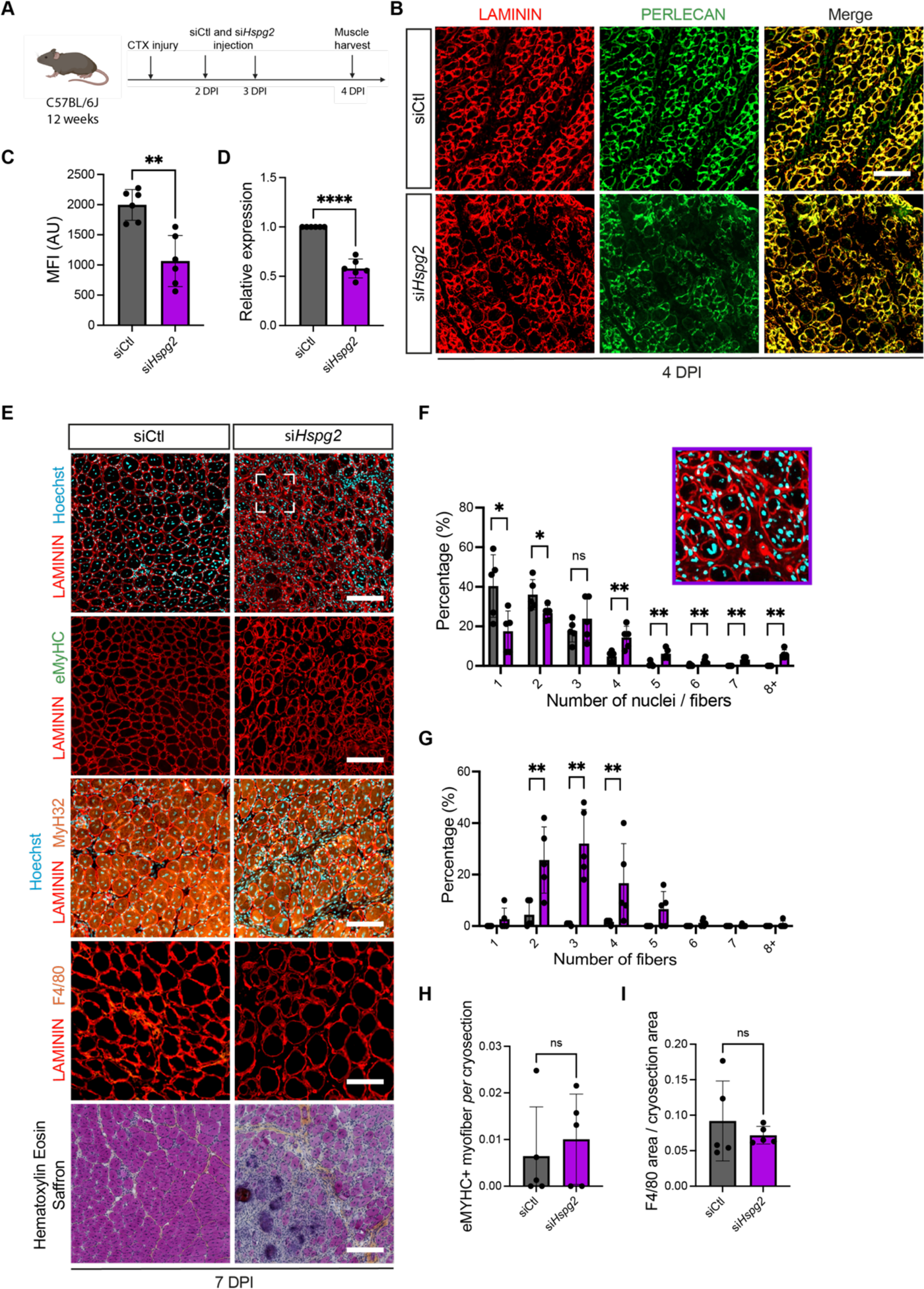
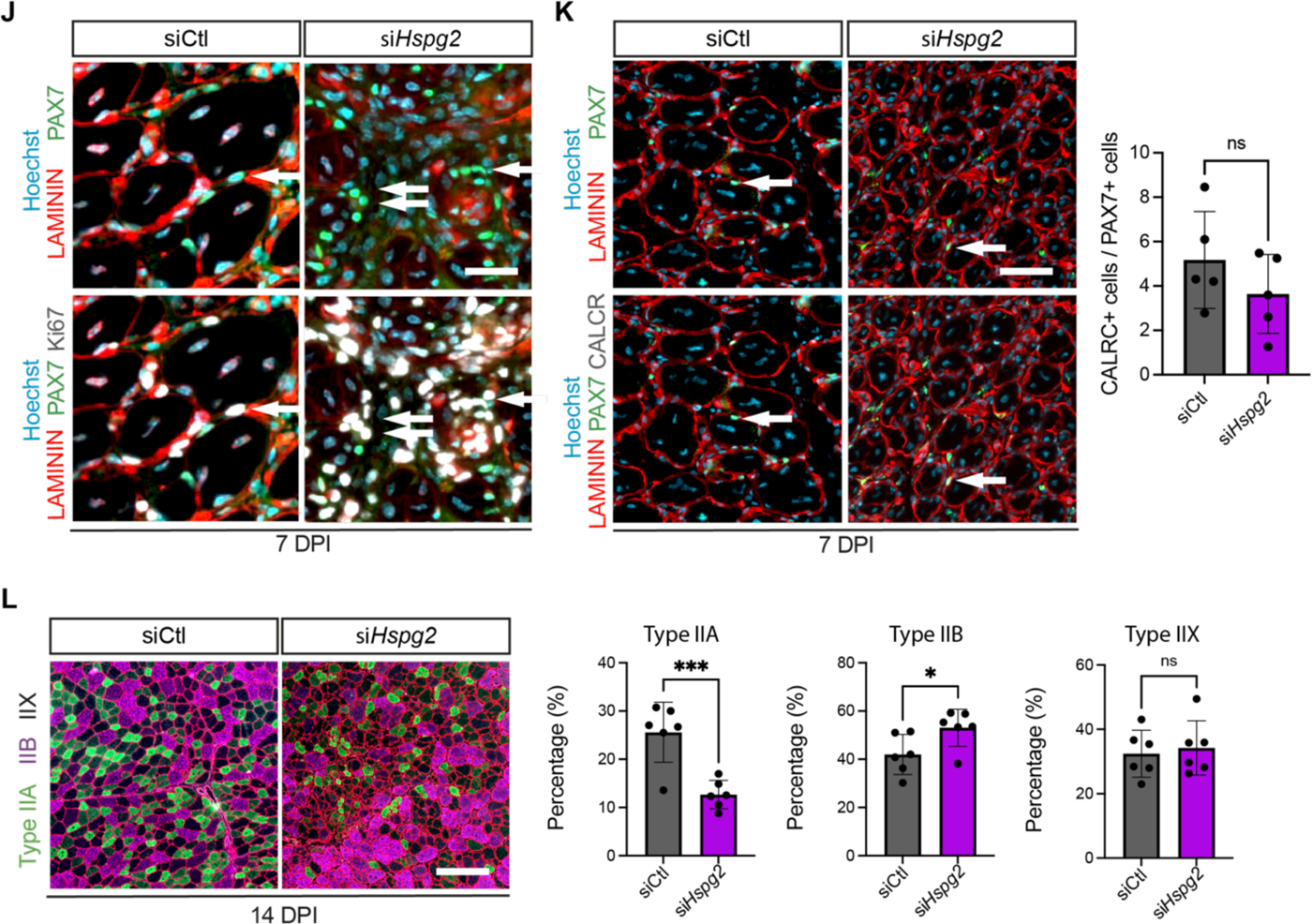
Characterization of muscle tissue phenotypes following si*Hspg2* knock-down. **(A)** Schematic of experimental timeline. **(B)** Perlecan and Laminin immunostaining in siCtl and siHspg2 muscles at 4 days post-injury (DPI). N=6 mice. **(C)** Mean Fluorescence Intensity of perlecan immunostaining in siCtl and siHspg2 treated muscles at 4 DPI. N=6 mice. **(D)** Relative expression of Hspg2 in 4 DPI siCtl and si*Hspg2* muscle cryosections. **(E)** Laminin, eMyHC, MY-32, F4/80 and Hematoxylin Eosin Saffron immunostaining of siCtl and si*Hspg2* transverse muscle sections at 7 DPI. N=5 mice. **(F)** Representative image and bar graph quantification of number of nuclei per fiber in siCtl and si*Hspg2* transverse muscle sections at 7 DPI. N=5 mice. **(G-I)** Bar graph quantification of **(G)** the number of fibers within a single basal lamina, **(H)** eMyHC+ myofibers, and **(I)** F4/80 area in siCtl and si*Hspg2* transverse muscle sections at 7 DPI. N=5 mice **(J)** Representative image of Pax7, Ki67 and Laminin immunostaining in siCtl and si*Hspg2* transverse muscle sections at 7 DPI. N=5 mice. **(K)** Representative Pax7, calcitonin receptor and laminin immunostaining (left) from siCtl and si*Hspg2* muscle transverse sections at 7 DPI and bar graph quantification (right) of the proportion of Pax7+ cells that are co-labeled for CalcrR+. N=5 mice. **(L)** Representative images of Type IIA and I myofiber staining in siCtl and si*Hspg2* transverse muscle sections at 14 DPI (left) and bar graph quantification (right) of the number of type IIA+, type IIB+ and type IIX+ myofibers. N=6 mice. P-values for *in vivo* data were calculated for Student t-test statistic, ***p˂0.001, *p˂0.05, ns: non significative. Scale bar, (B) 50 μm, (E) 75 μm, 75 μm, 75 μm, 50 μm, 100 μm, (J) 25 μm, (K) 50 μm, (L) 200 μm

**Figure S8.**
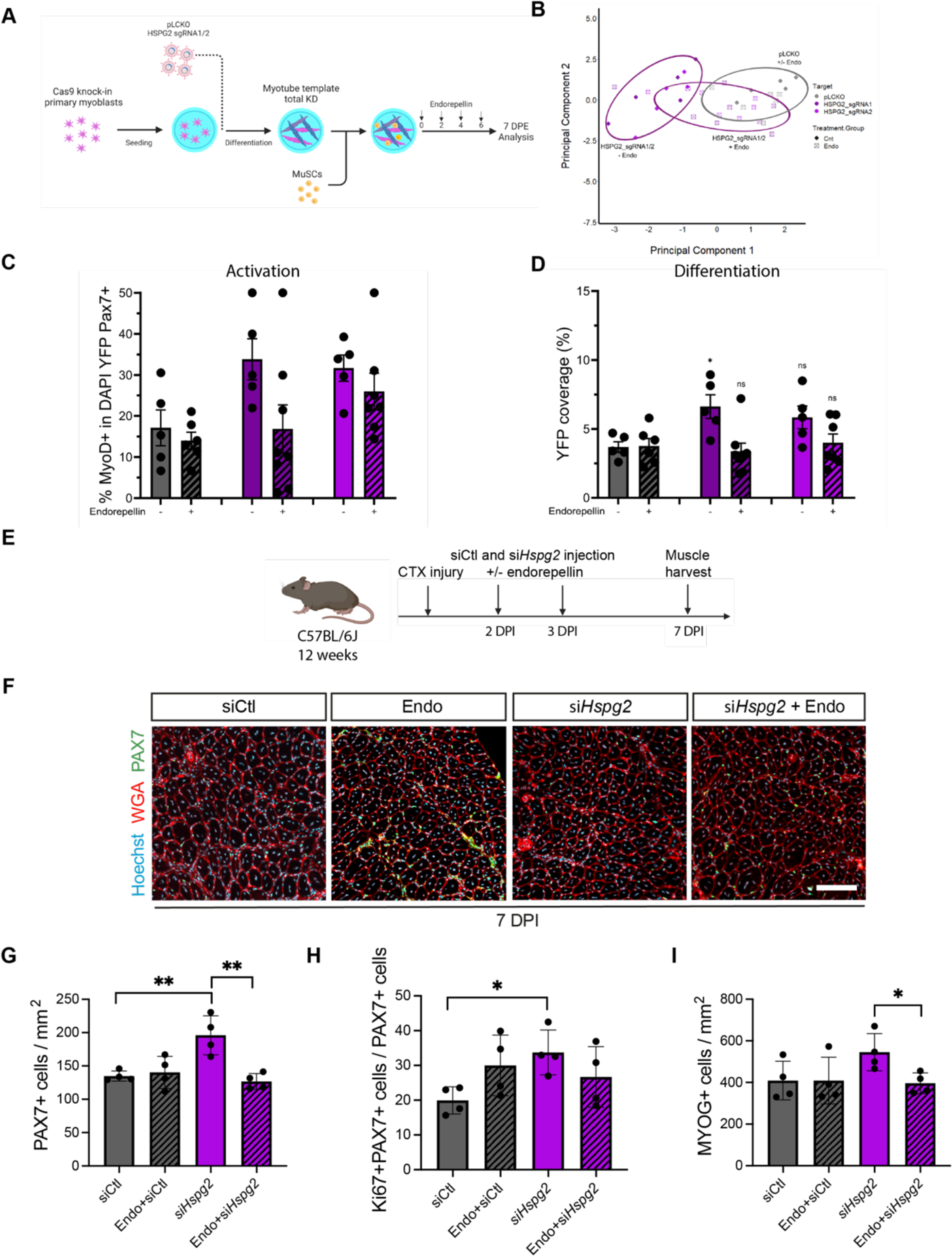
Hspg2 knock-down is partially rescued by endorepellin in mini-IDLE. **(A)** Schematic representation of the experimental setup for the testing of endorepellin on MuSCs in the mini-IDLE assay (made with BioRender). **(B-C)** Bar plots showing the percentage of Pax7+EdU- nuclei (DAPI+) **(B)** and cells per mm^2^ (DAPI+) **(C)** on day 3 across control and different concentrations (per mL) of endorepellin. Graphs display mean ± s.e.m. with individual technical replicates. one-way ANOVA with Dunnet’s test for each experimental condition compared against the control. n=4-5 wells across N=2 independent biological replicates. *p˂0.05, ***p˂0.001 (B); *p˂0.05 (C). **(D-H)** Bar plot showing mononuclear DAPI YFP Pax7+ cell density **(D)** percentage MyoD+ in the mononuclear DAPI YFP Pax7+ cell population **(E)** average max/min feret diameter of the mononuclear DAPI YFP Pax7+ cell population **(F)** average nuclear eccentricity of the mononuclear DAPI YFP Pax7+ cell population **(G)** and average YFP coverage per FOV **(H)** across transduction with different sgRNAs (color coded) with or without endorepellin (x-axis). Graphs display mean ± s.e.m. with individual technical replicates; one-way ANOVA with Dunnet’s test for each experimental condition compared against the control (pLCKO - endorepellin). n=5-8 tissues across N=2-3 independent biological replicates. ns (D); ns (E); **p=0.0017, 0036 ***p=0.0002, 0.0004 (F); *p=0.0332, 0.0279 (G); *p=0.0283 (H).

**Figure S9.**
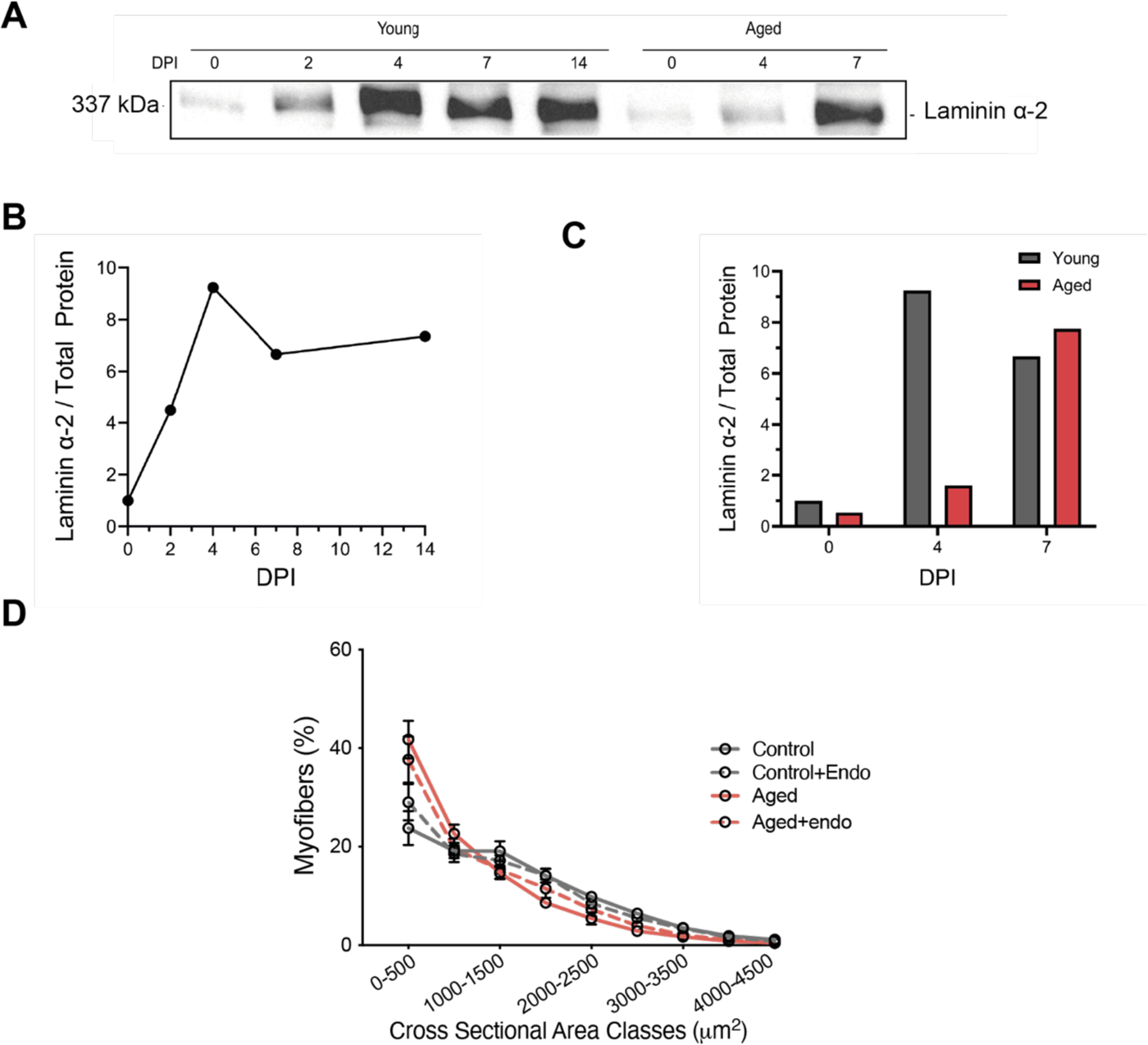
Basal lamina and myofiber reconstruction in young and aged regenerating mouse muscle. **(A)** Representative western blot for laminin α-2 in young vs aged mouse muscle lysate across days post-injury (DPI). **(B)** Laminin α-2 protein quantification in young mice across two weeks post-injury. **(C)** Bar plot showing laminin α-2 levels in the young (red) and aged (grey) mouse lysate at 0, 4 and 7 DPI. Plot displays mean. **(D)** Line plot showing myofiber size distribution (cross-sectional area) at 14 DPI in young (control, grey) and age (red) mouse muscle transverse sections without (solid line) and with added endorepellin (dashed lines). N=4-5 mice. Plot displays mean ± s.d.

